# Genetic, neural and early-environmental architecture of resilience

**DOI:** 10.64898/2026.11.16.767004

**Authors:** Tung-Yen Wu, Yun-Yen Yang, Yu-Jen Liang, Fa-Hsuan Lin, Hsin-Chou Yang, Chen-Ying Huang

## Abstract

Resilience is shaped by factors ranging from the genome through early-life experience to the brain, yet its biological basis in humans remains unclear because studies have rarely integrated these levels and have been confined to small samples. Using the Brief Resilience Scale (BRS) and multimodal data from tens of thousands of UK Biobank (UKB) participants, with replication in Generation Scotland (GS), we found that resting-state functional connectivity, particularly within the flexibility and valuation networks, is a robust neural correlate of resilience. Resilience also showed a generalizable genetic basis: a polygenic score (PGS) derived from a UKB genome-wide association study predicted BRS in GS. Genes and childhood experiences converged on the flexibility network through distinct routes: the polygenic association with resilience was partly explained by central executive–salience integration, whereas childhood experiences were associated with default-mode-centered integration. By separating genetically anchored from experience-sensitive routes, these findings point to network targets that interventions could tailor to the sources of an individual’s vulnerability.

## Introduction

Humans adapt to stress heterogeneously; some experience persistent psychological difficulties, while others quickly recover through resilience. Resilience refers to one’s ability to bounce back from stressful events (Fig. 1a). Factors that contribute to successful bounce back and adaption are termed resilience factors (RFs)^1,2^. These RFs can be broadly classified into biological, psychological, social, and ecological domains, which we collectively conceptualized as biopsychosocial-ecological RFs^3,4^ (Fig. 1b; left). Increasing attention has focused on identifying biological RFs, as doing so could help stratify individual resilience by recognizing fixed predispositions and provide malleable targets for clinical intervention^1–7^. To this end, large-scale approaches, such as genome-wide association studies (GWAS) and brain-wide association studies (BWAS), offer promising avenues for systematically uncovering biological signatures underlying resilience (Fig. 1b; right).

**Fig. 1.**
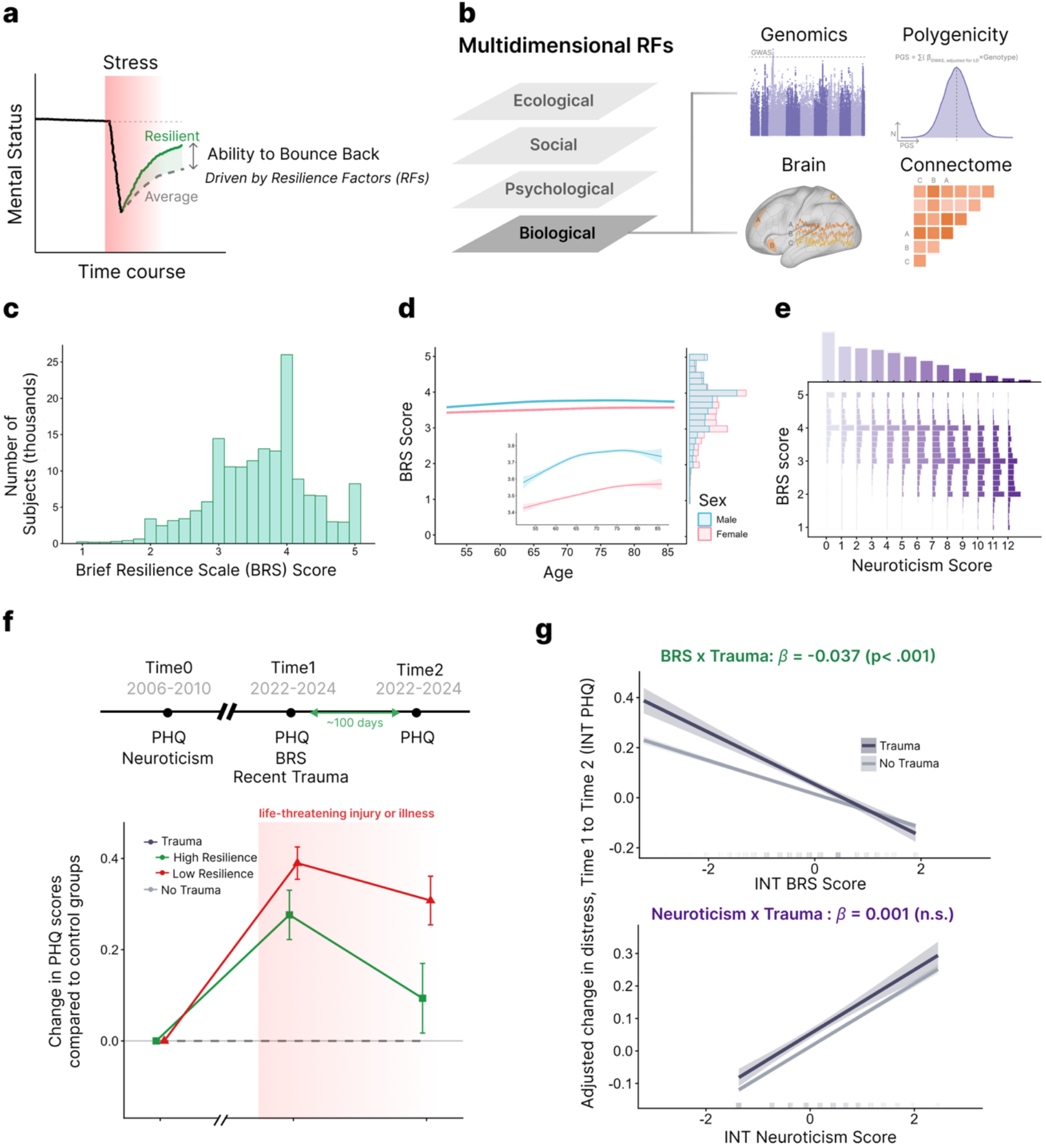
Conceptual framework and empirical validation of the Brief Resilience Scale (BRS). **a**, Schematic of resilience as the capacity to recover mental status following stress exposure. **b**, Multidimensional resilience factors (RFs), illustrating the study’s focus on genetic and brain-related biological RFs. **c, d**, Phenotypic characterization of BRS in the UK Biobank (UKB) GWAS discovery cohort (N = 159,575): score distribution (c) and association with age and sex (d; inset, zoomed view of trajectories; shaded bands, 95% CI). **e–g**, Analyses in the BRS validation cohort (N = 110,346). **e**, Joint distribution of BRS and neuroticism scores. **f**, Top, study design across three waves (Time 0–2). Bottom, excess PHQ change from Time 0 after life-threatening injury or illness, by BRS level (median split), relative to BRS-matched unexposed participants, with entropy balancing on Time 0 PHQ (N = 105,359). Error bars, bootstrap s.e. **g**, Predicted change in distress from Time 1 to Time 2 (INT PHQ) as a function of INT BRS (top) and neuroticism (bottom) scores, by trauma exposure. Shaded bands, 95% CI.

Despite its immense societal and clinical importance, elucidating the biological architecture of resilience remains a formidable challenge due to the dilemma of phenotyping resilient subjects. A well-accepted consensus defines resilience as maintaining better-than-expected mental health following adversity^1^; however, such outcome-based measurements require specific trauma exposure, severely restricting sample availability^5^. This limitation poses a major bottleneck, as sample sizes in the thousands are typically required to achieve sufficient statistical power in GWAS^8^ and BWAS^9^. Conversely, while self-report psychometric scales offer a scalable proxy for large-scale cohorts, skepticism persists regarding whether questionnaire-derived scores genuinely capture dynamic behavioral resilience^1,10,11^. Moreover, these scores overlap substantially with negative personality traits such as neuroticism, both phenotypically^12^ and genetically^13^. Whether questionnaire-based resilience can nonetheless serve as a biologically meaningful phenotype is therefore a pressing question.

Owing to these phenotyping constraints, the genetic and neural architectures underlying resilience remain poorly understood. At the genetic level, although resilience is moderately heritable, no genome-wide significant loci identified in prior GWAS have been successfully replicated across independent cohorts^13^, casting doubt on the existence of a generalizable genetic architecture. At the neural level, candidate brain network hypotheses have largely been guided by either psychological RFs, including positive appraisal style^2^ and regulatory flexibility^2,14^, or affect-based resilience mechanisms (RMs), such as down-regulating negative and up-regulating positive affect^15^. Together, these theoretical frameworks have converged onto three commonly reported hypotheses regarding resilience-related intrinsic brain networks ((1) the triple network, (2) the prefrontal-amygdala network, and (3) the prefrontal-striatum network. However, while resting-state functional connectivity (rsFC) provides a powerful window into the intrinsic functional architecture supporting these processes, prior rsFC studies of resilience have been predominantly under sampled, yielding contradictory findings with opposite effect directions. For example, stronger connectivity between the anterior insula and the medial temporal lobe has been reported to be associated with greater resilience in both outcome-based^16^ and questionnaire-based empirical studies^17^, yet another outcome-based study reported elevated connectivity between these regions in less-resilient individuals^18^, with each direction interpreted under incompatible theoretical accounts. Well-powered rsFC evidence is therefore needed to determine which of these candidate networks robustly track resilience. Moreover, because genetic and neuroimaging investigations have largely operated in isolation, it also remains unclear how these candidate brain networks bridge fixed predispositions and modifiable environmental RFs to shape resilience.

Here, we address these phenotyping, genetic and neuroimaging bottlenecks with a gene–brain–behavior framework spanning the UK Biobank (UKB) and the independent Generation Scotland (GS) cohort. We begin by showing that the Brief Resilience Scale (BRS) captures recovery after adversity beyond neuroticism. With a phenotype that is both valid and available at biobank scale, we can then examine the biology of resilience with the statistical power such studies require. A linear mixed-model GWAS in UKB yields a polygenic score (PGS) that, for the first time, predicts resilience in an independent cohort, and its functional and cellular annotations point to neuronal communication and glutamatergic neurons. Building on this convergence on neural signaling, we turn to intrinsic brain networks: in the largest resilience rsFC study to date, we jointly test the triple, cortical–amygdala and cortical–striatal network hypotheses, of which the triple and cortical–striatal networks replicate in GS. Having identified network features that may carry fixed genetic predisposition, we next ask how they relate to a modifiable factor, early-life environment, and whether the two converge on the same circuits. Integrating PGS, childhood experiences, rsFC and BRS, we find that polygenic predisposition and early-life environment map onto partly distinct intrinsic brain correlates of resilience.

## Results

### Cohort overview and phenotypic characterization of BRS

In this study, the primary GWAS discovery cohort comprised 159,575 participants in UKB (Fig. S1b; see Methods). For the behavioral-level investigation, this GWAS discovery cohort was filtered by the completeness of longitudinal mental health and adversity records, yielding a subset of 110,346 individuals (Fig. S1b). In parallel, to enable unbiased imaging-genetics analyses, the same GWAS discovery cohort (N = 159,575) was partitioned into a non-overlapping GWAS for PGS cohort without rsfMRI data (N = 130,726) and an imaging-genetics discovery cohort with complete functional connectivity matrices (N = 27,729, after excluding 1,120 individuals with incomplete matrices; Fig. S1b).

In the independent GS replication dataset, a total of 8,702 participants had complete genotype, BRS, and covariate records, forming the replication cohort for PGS validation (Fig. S1c). A neuroimaging subset of 1,060 participants with post-QC rsfMRI data was used to replicate imaging-related discovery in UKB (Fig. S1c). Among them, 997 subjects had complete genotypes and were used to replicate the imaging-genetics-related discovery in UKB (Fig. S1c).

The following paragraph characterizes the phenotypic profile of BRS in the UKB GWAS discovery cohort (N = 159,575). BRS spanned the full-scale range (1–5), with a mean of 3.61 (SD = 0.74; median = 3.67; Fig. 1c). Scores showed a mild negative skew (skewness = −0.32; IQR = 3.17–4.00), which was normalized via rank-based inverse normal transformation (INT) for subsequent genetic and neuroimaging analyses. Demographic analyses revealed linear and non-linear associations with age (age: β = 0.0047 per year, 95% CI [0.0042, 0.0052], P = 5.1 × 10⁻⁷⁸; centered age-squared: β = −0.00023, 95% CI [−0.00029, −0.00017], P = 4.3 × 10⁻¹³) and significant sex differences, with higher scores in males (mean 3.74 vs 3.52 in females; adjusted β = −0.21, 95% CI −0.22 to −0.21, P < 1 × 10⁻³⁰⁰; Fig. 1d). These associations motivated the inclusion of age and sex as covariates in all subsequent genetic and neuroimaging models. To evaluate construct validity, we tested the relationship between BRS and neuroticism in the BRS validation cohort (n = 110,346) and observed a substantial negative correlation (r = −0.46, 95% CI [−0.47, −0.46], P < 1 × 10⁻³⁰⁰; Fig. 1e). Thus, although higher resilience was strongly coupled with lower negative affectivity, the two traits shared only about 21% of their variance, indicating that BRS retains distinct phenotypic variance beyond the mere absence of neuroticism.

### BRS predicts resilient outcome

The goal of this study was to characterize the genetic and neural correlates of psychological resilience, defined as the capacity to recover following adversity. As a foundation for these analyses, we first examined whether scores on the BRS predict recovery beyond their association with general mental health. Using trauma events and longitudinal mental health records in UKB, we compared changes in mental health between trauma-exposed and unexposed individuals at the same BRS level (Fig. 1f). The unexposed group served as a reference at each BRS level, allowing us to distinguish better mental health in general from better recovery following adversity. If BRS captures the capacity to bounce back, trauma-exposed individuals with higher scores should return closer to the mental health levels of their unexposed counterparts during follow-up. Consistent with this prediction, individuals with higher BRS scores showed a closer return to their unexposed counterparts following life-threatening injury or illness (Fig. 1f): When both BRS groups showed similar initial increases in PHQ scores, the high-BRS group had approached its unexposed reference by Time 2 (excess PHQ change from Time 0: 0.09 points, 95% CI [−0.06, 0.24], P = 0.22). In contrast, PHQ scores in the low-BRS group remained elevated relative to its unexposed reference (0.31 points, 95% CI [0.20, 0.41], P = 7.6 × 10⁻⁹).

Within this trauma-specific recovery framework, we tested the interaction between BRS and trauma exposure on symptom change from Time 1 to Time 2 (N = 110,346; trauma-exposed n = 8,319). The model adjusted for pre- and post-event PHQ scores, neuroticism (and its interaction with trauma), age and sex. Trauma exposure was associated with poorer recovery over this interval (β = 0.039, 95% CI [0.025, 0.054], P = 1.0 × 10⁻⁷), but this association weakened with increasing BRS (BRS × trauma interaction: β = −0.037, 95% CI [−0.053, −0.021], P = 5.0 × 10⁻⁶; Fig. 1g top). Because the main and interaction effects were of comparable magnitude but opposite sign, a 1 S.D. increase in BRS was sufficient to offset the average impact of trauma. Consistently, simple-slope analysis showed that the effect of trauma was largest at 1 S.D. below the mean BRS (β = 0.076, 95% CI [0.055, 0.097], P = 6.3 × 10⁻¹³) and was not distinguishable from zero at 1 S.D. above the mean (β = 0.002, 95% CI −0.020 to 0.025, P = 0.83). These results suggest that, at BRS scores approximately 1 S.D. above the population mean, trauma-exposed and unexposed individuals showed no detectable difference in PHQ change (Fig. 1g top).

To assess whether the association between BRS and recovery extended beyond general emotional vulnerability, we examined the role of neuroticism. The result showed that its interaction with trauma was not significant when the BRS × trauma interaction was included (β = 0.001, 95% CI [−0.0, 0.018], P = 0.89). When the BRS × trauma term was omitted, the neuroticism × trauma interaction was significant (β = 0.019, 95% CI [0.005, 0.033], P = 0.008). Given the shared variance between the two traits (r = −0.46), the neuroticism × trauma interaction appeared to depend on variation it shares with BRS, whereas the BRS × trauma interaction was independent of neuroticism.

Because trauma-exposed participants had higher pre-exposure distress than unexposed participants (standardized mean difference for Time 0 PHQ = 0.46; Fig. S2a), we conducted two sensitivity analyses to examine whether this baseline difference affected the BRS × trauma interaction: entropy balancing on Time 0 PHQ, and stratification by Time 0 PHQ severity. After entropy balancing, the interaction was attenuated but remained significant (β = −0.027, 95% CI [−0.043, −0.011], P = 8.9 × 10⁻⁴; Fig. S2b). In the stratified analysis, the interaction did not differ across severity strata (β range −0.045 to −0.030; heterogeneity F(2) = 0.28, P = 0.76; Fig. S2c). Moreover, both the unexposed reference and the baseline adjustment were needed to avoid inflating the estimate (Fig. S2d): without adjustment for Time 0 PHQ, the BRS × trauma interaction was 45% larger (β = −0.053, 95% CI [−0.070, −0.037]). Among trauma-exposed participants alone, the BRS slope was about three times larger (β = −0.106, 95% CI [−0.123, −0.089]), and four times larger when Time 0 PHQ was also omitted (β = −0.150, 95% CI [−0.168, −0.133]), because this slope also includes the association between BRS and symptom change that is shared with unexposed participants (β = −0.067). These findings suggest that BRS captures variation in recovery following adversity beyond general differences in mental health and emotional vulnerability.

### Genetic factors and cell types through genome sequencing data

Building on longitudinal evidence linking BRS to recovery following adversity, we next examined its genetic architecture. We conducted a GWAS of BRS in UKB using a linear mixed model (BOLT-LMM; N = 159,575). The SNP-based heritability is 7.32% by LDSC estimation. To validate the polygenicity of BRS measurements, we used the GS cohort’s genotypic data to compute the PGS by LDpred2 (*N* = 8,702). The calculated PGS showed a robust effect size on BRS (Fig. 2b), where a one standard deviation increase in the PGS was associated with a β of 0.080 (95% CI [0.059, 0.101]; *P* = 6.7 × 10⁻¹⁴), after adjusting for sex, age and centered age squared. This PGS was derived with the infinitesimal model of LDpred2 (LDpred2-inf). We chose this model after comparing it with the automatic model (two models performed similarly, but infinitesimal provides the more reproducible basis for the future analyses; details in the Supplementary Information and Fig. S3).

**Fig. 2.**
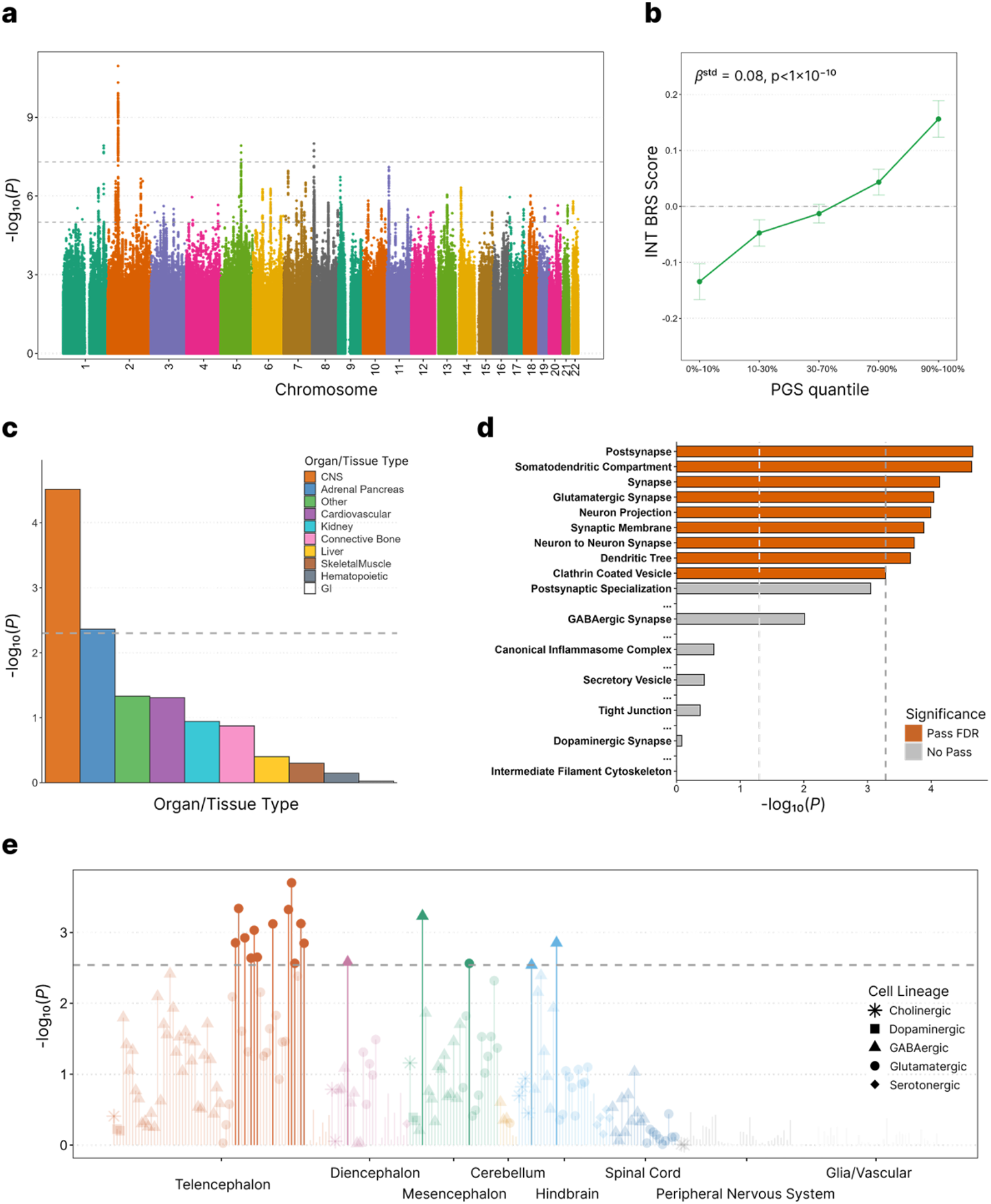
Genetic architecture and cellular enrichment of BRS. **a,** Manhattan plot of the linear mixed-model (BOLT-LMM) genome-wide association study (GWAS) for Brief Resilience Scale (BRS) scores in the UK Biobank discovery cohort (N = 159,575). The dashed horizontal line indicates the genome-wide significance threshold (P = 5 × 10⁻⁸ and P = 1 × 10⁻^5^). **b,** Validation of the polygenic score (PGS) in Generation Scotland (GS; N = 8,702), derived via LDpred2 using GWAS weights from a (error bars, s.e.m.). **c,** Partitioned heritability enrichment across 10 human tissue/organ groups estimated via linkage disequilibrium score regression (LDSC). **d,** Over-representation analysis of MAGMA-prioritized genes across selected Gene Ontology Cellular Component (GO-CC) terms. **e,** Cell-type-specific enrichment across nervous system regions (colors) and neurotransmitter lineages (shapes) in the Linnarsson mouse brain single-cell atlas (Dashed lines in c–e, PFDR < 0.05; light dashed line in d, nominal P = 0.05).

To further translate this genetic variation to candidate mechanisms underlying individual differences in resilience, we next performed the enrichment analysis to identify the tissue and cellular components implicated by these genetic associations. By leveraging the previous annotation work mapping SNP-level chromatin marks across 10 human tissue groups, we found significant enrichment of BRS heritability in the central nervous system (Fig. 2c), with possible additional enrichment in the adrenal/pancreas group. We next examined cellular components using gene-level overrepresentation analysis based on the Gene Ontology Cellular Component (GO-CC) dataset. This analysis showed that 9 components were significantly over-represented (FDR < 0.05) out of 642 tested components. These leading components are predominantly synaptic and neuronal compartments (Fig. 2d). Cellular components related to frequently hypothesized resilience factors, such as blood-brain barrier integrity (e.g., tight junctions) and neuroinflammation (e.g., canonical inflammasome complex), were not significantly enriched in our genetic data. Among neurotransmitter systems, the enrichment was most pronounced for glutamatergic synapses, with marginal evidence for GABAergic involvement and no association for dopaminergic synapses. While GO-CC relies on a priori curated gene sets, we sought to confirm these findings using empirical gene expression data. We integrated single-cell gene expression data from the Linnarsson mouse nervous system atlas and the Allen human brain atlas using FUMA cell-type analysis, and grouped cell types by their principal neurotransmitter. In the mouse atlas, 17 of 265 cell types were significantly enriched (FDR < 0.05; Fig. 2e). Glutamatergic neurons showed the most robust enrichment, accounting for 13 of the 17 significant cell types (13 of 65 glutamatergic types), including 12 of the 24 telencephalic excitatory projection neuron types (strongest association, P = 2.0 × 10⁻⁴, FDR = 0.033). GABAergic neurons showed weaker and more restricted signals: 4 of 79 GABAergic types reached significance, none of which were among the 27 telencephalic GABAergic types (21 interneuron and 6 medium spiny neuron types). Dopaminergic neurons remained non-significant (0 of 4; all P > 0.40), as did serotonergic, cholinergic and non-neuronal cell types. In the human brain atlas, the pattern was similar: the three cell types reaching nominal significance were all glutamatergic neurons of the middle temporal gyrus (MTG), and nine of the ten top-ranked MTG cell types were glutamatergic (Fig. S4). Together, these findings implicate forebrain glutamatergic neurons and their synapses as a likely cellular substrate of BRS genetics, motivating our subsequent examination of how this genetic variation is expressed at the level of large-scale brain networks.

### Brain networks assessed by resting-state fMRI

Three commonly hypothesized brain systems for RFs include: (H1) the Large-Scale Flexibility Network, in which the salience network (SN) coordinates switching between the default mode (DMN) and central executive networks (CEN) to support cognitive flexibility; (H2) the Cortico-Amygdala Regulatory Network, in which prefrontal regulation of the amygdala supports reappraisal and emotion regulation; and (H3) the Cortico-Striatal Valuation Network, in which ventral striatal circuitry drives active coping. To facilitate comparison with previous and future studies, we defined a consistent anatomical framework for defining the networks: A priori regions of interest (ROIs) as network nodes were selected based on predefined parcellations (the Glasser cortical and Tian subcortical atlases) and previous literature (Fig. 3a; see Supplementary Information). We calculated ROI-to-ROI resting-state functional connectivity (rsFC) for 45, 17, and 57 pairwise connections (edges) in H1, H2, and H3, respectively. In UKB (N = 27,729), we tested associations between each edge and BRS using partial correlations adjusted for age, mean-centered age squared, sex, log head motion, and log tSNR. Significant associations were observed for all 45 edges in H1, 9 of 17 in H2, and 50 of 57 in H3 (FDR-adjusted P < 0.05), with partial r ranging from 0.012 to 0.052 (Fig. 3b). We next assessed whether the three hypothesized systems made distinct contributions to explaining variation in BRS. Using nested partial F-tests, we tested whether each system explained additional variance after accounting for the other two. Each system explained unique variance in BRS beyond the other two (H1: F(45, 27,604) = 3.23, P = 1.9 × 10⁻¹²; H2: F(17, 27,604) = 2.64, P = 2.5 × 10⁻⁴; H3: F(57, 27,604) = 1.80, P = 2.1 × 10⁻⁴; all below the Bonferroni threshold; Fig. 3c), with the strongest evidence for the Large-Scale Flexibility Network.

**Fig. 3.**
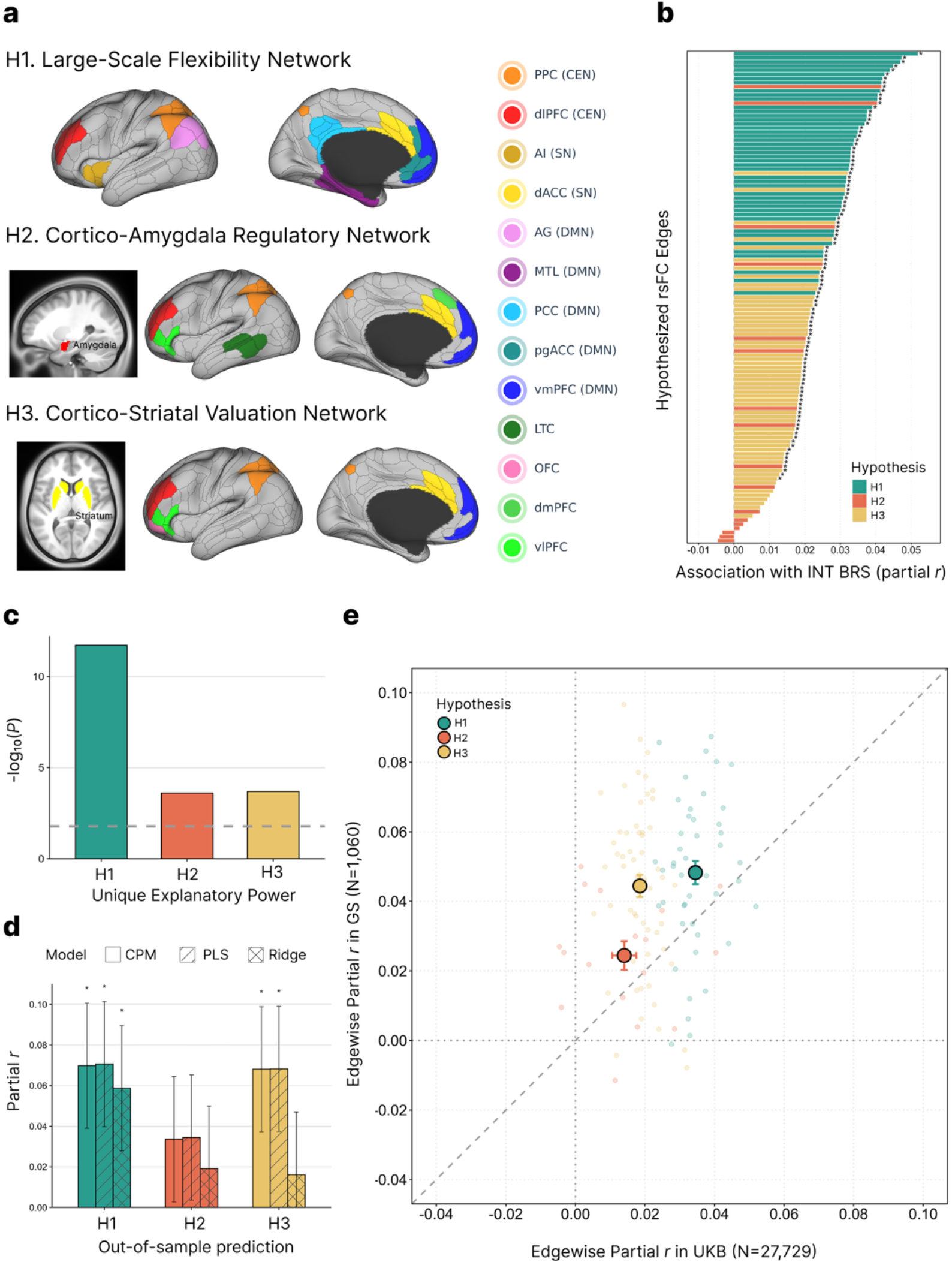
Intrinsic brain network correlates of BRS. **a,** Anatomical ROIs composing the three hypothesized networks, defined from the Glasser cortical parcellation and the Tian subcortical atlas. H1, the large-scale flexibility network, comprises the central executive network (CEN; dlPFC, PPC), salience network (SN; AI, dACC) and default mode network (DMN; AG, MTL, PCC, pgACC, vmPFC). H2, the cortico-amygdala regulatory network, links the amygdala with prefrontal, parietal and temporal regions. H3, the cortico-striatal valuation network, links the striatum with prefrontal and parietal regions. Colors denote ROIs. **b,** Edge-wise partial correlations between each of the 119 ROI-level functional connections (FCs) and INT BRS scores in the UKB imaging-genetics discovery cohort (N = 27,729), adjusted for age, mean-centered age squared, sex, log head motion and log tSNR. FCs are ordered by effect size and colored by hypothesis; stars mark FCs with FDR significance. **c,** Unique explanatory power of each network, assessed by partial F-tests comparing a full model containing all 119 FCs with a reduced model omitting the target network. The dashed line indicates the Bonferroni-corrected threshold for three networks (P = 0.05/3). **d,** Out-of-sample prediction in the GS imaging replication cohort (N = 1,060) using network-specific models trained in UKB (CPM, PLS, and Ridge). Predictive power is the partial correlation between the predicted score and INT BRS, adjusted for age, age squared, sex, imaging site, log head motion and log tSNR. Error bars, ±1 s.e.; stars, one-tailed P < 0.05. **e,** Edge-wise partial correlations in UKB (x axis) versus GS (y axis). Small points represent individual FCs; large points represent the network mean (error bars, s.e.m. across FCs on both axes). The dashed line marks identity.

We next tested replication in GS (N = 1,060), whose fMRI processing pipeline was harmonized with the UKB protocol (see Supplementary Information). Network-specific models trained in UKB were applied to GS. For H1, all three models significantly predicted BRS (Fig. 3d; CPM, r = 0.070, P = 0.012; PLS, r = 0.071, P = 0.011; Ridge, r = 0.059, P = 0.028; one-tailed). H3 was replicated by CPM (r = 0.068, P = 0.014) and PLS (r = 0.068, P = 0.013) but not by Ridge (r = 0.016, P = 0.30), whereas H2 did not replicate (r ≤ 0.034, P ≥ 0.13). These predictive correlations exceeded the largest single-edge effects in UKB (r = 0.052), consistent with many weak, same-signed FC effects that are individually unreliable but add up to a stronger and more stable signal when aggregated at the network level (see Supplementary Information and Fig. S5 for a comparison of predictive models). Individual edge associations had consistent directions across cohorts (Fig. 3e). GS estimates matched the UKB direction for 44 of 45 edges in H1, 13 of 17 in H2, and 54 of 57 in H3. We evaluated two measures: the number of edges with matching directions and the mean GS partial correlation after aligning each edge to its UKB direction. To account for dependence among edges, we used permutation tests that shuffled BRS within each GS imaging site while keeping connectivity data fixed (see Supplementary Information for details). Directional concordance exceeded chance for H1 (P = 0.042) and H3 (P = 0.029; Fig. S6a). The mean sign-aligned partial correlations also exceeded the permutation null for H1 (mean r = 0.048, P = 0.011) and H3 (mean r = 0.044, P = 0.009; Fig. S6b). Neither measure was significant for H2 (P ≥ 0.11). Together with the prediction results, these findings support cross-cohort consistency in the associations of H1 and H3 connectivity with BRS. Notably, the associations were positive across H1: edges within and between the CEN, SN and DMN were positively associated with BRS for all 45 edges in UKB and 44 of 45 in GS (Fig. S6c). This pattern only partly agrees with previous empirical studies and reviews, as addressed in the Discussion.

### Brain network mediation of genetic resilience factors

To delve deeper into the neural mechanisms linking our polygenic findings to neuronal communication, we asked whether rsFC could account for part of the association between PGS and BRS. This required a PGS whose genetic weights were estimated without including the participants used for imaging analyses. We therefore divided the UKB GWAS sample into participants with and without MRI data (Fig. S1b). We repeated the GWAS in participants without MRI data (N = 130,726) and used the resulting summary statistics to calculate PGS in the imaging sample (N = 27,729). The derived PGS was robustly associated with BRS in both the UKB imaging sample (N = 27,729; β = 0.067, P = 1.4 × 10⁻²⁸) and GS (N = 8,702; β = 0.077, P = 5.7 × 10⁻¹³), supporting its use in subsequent connectivity analyses.

We first examined whether PGS was associated with rsFC. For illustration, we plotted two prefrontal connections within H1, dlPFC–dACC and vmPFC–dACC, across PGS quartiles in UKB (N = 27,729; Fig. 4a). Higher PGS was associated with stronger dlPFC–dACC connectivity (β = 0.019, P = 0.001), whereas the association with vmPFC–dACC connectivity was not significant (β = 0.005, P = 0.40). We then conducted a separate mediation analysis for each edge to test whether polygenic variability manifests as psychological resilience through specific rsFC edges (Fig. 4b). Significant indirect effects were observed for 13 of 45 edges in H1, with none reaching significance in H2 [0 of 17] or H3 [0 of 57]; Fig. 4c). Standardized indirect effects (a × b) across these 13 edges ranged from 3.2 × 10⁻⁴ to 8.2 × 10⁻⁴ (all FDR < 0.05). Nine of these connections were within or between the CEN and the SN. To estimate their combined contribution, we entered the 13 edges together in a parallel multiple mediator model. Their total standardized indirect effect was 1.1 × 10⁻³ (95% CI [0.37, 1.87] × 10⁻³), accounting for 1.6% of the PGS–BRS association (95% CI [0.5, 2.8] %). Together, these findings support a small indirect association between PGS and BRS through selected H1 connections, with most significant connections involving the CEN and SN.

**Fig. 4.**
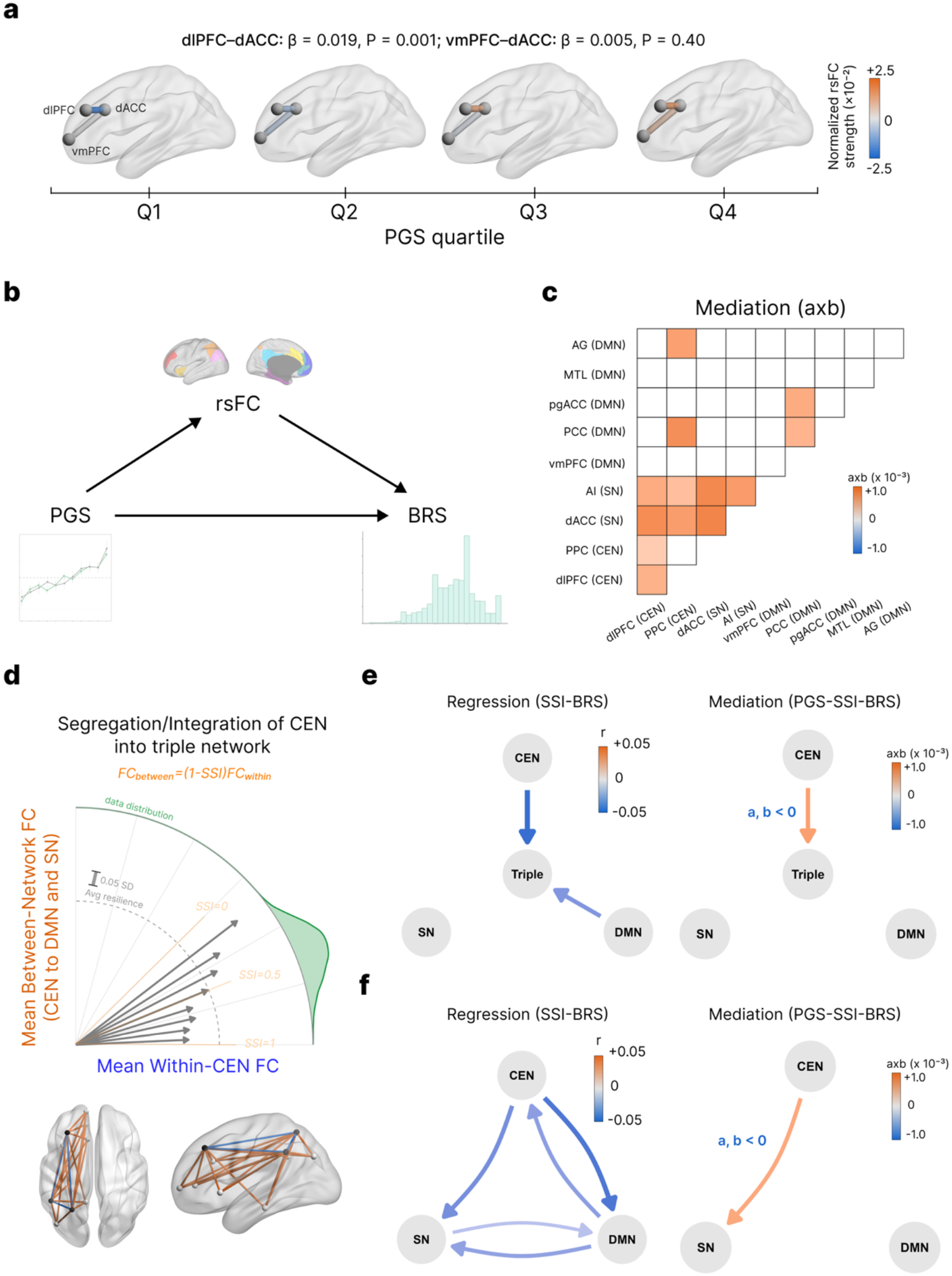
Edge- and system-level connectomic mediation of polygenic resilience. **a,** Functional connectivity (FC) of dlPFC–dACC and vmPFC–dACC across quartiles of the resilience polygenic score (PGS) in the UKB imaging-genetics discovery cohort (N = 27,729). Edge color indicates the mean normalized rsFC. Associations between the PGS and each FC (standardized β), adjusted for covariates, are shown above. **b,** Mediation model. The PGS (X) is associated with INT BRS (Y) through FC (M). **c,** Edge-wise completely standardized indirect effects (a × b) of the 45 FCs in the large-scale flexibility network (H1) in UKB (N = 27,729), adjusted for the covariates. Only FCs with FDR-significant indirect effects are colored. **d,** Schematic of the segregation–integration index (SSI) of the CEN within the triple network. For each participant, within-CEN FC (x axis) and mean between-network FC of the CEN with the SN and DMN (y axis) define an angle; dashed lines mark SSI = 1, 0.5 and 0. Arrows represent participants binned by angle (5° bins), with arrow length indicating the mean INT BRS in each bin relative to the sample average (dashed arc); the green density shows the distribution of participants (UKB imaging-genetics discovery cohort, N = 27,729). Brain renderings illustrate within-CEN (blue) and between-network (orange) connections. **e,** One-to-triple SSI, quantifying the segregation of each network from the other two networks of the triple network. Left, partial correlations between SSI and INT BRS; right, standardized indirect of the PGS on INT BRS through SSI. **f,** Pairwise SSI, in which an arrow from network X to network Y denotes the segregation of X from Y; left and right panels as in **e**. In **e** and **f**, SSIs were rank-based inverse normal transformed, models were adjusted for the covariates and only significant paths are shown.

To assess whether this pattern extended beyond UKB, we repeated the mediation analyses in GS cohort (N = 997). H1 indirect effects matched the UKB direction for 42 of 45 edges (permutation P = 0.047; Fig. S7a). The mean GS indirect effect, after aligning each edge to its UKB direction, also exceeded the permutation null (mean standardized indirect effect = 2.1 × 10⁻³, permutation P = 0.022; Fig. S7b). Neither measure was significant for H2 (directional concordance: 12 of 17, P = 0.072; sign-aligned mean: P = 0.070) or H3 (directional concordance: 27 of 57, P = 0.55; sign-aligned mean: P = 0.57). Although no individual FC reached FDR significance in GS, the UKB estimates fell within the GS 95% confidence intervals, and the GS estimates lay within the range expected for a cohort of this size (Fig. S7c; Supplementary Information for details).

### Relative balance between brain networks as resilience factors

The edge-level mediation findings implicated specific H1 connections, particularly those within or between the CEN and SN (Fig. 4c). Under the large-scale flexibility hypothesis, resilience may depend partly on coordination among the CEN, SN, and DMN. We therefore extended the analysis from individual connections to the relative integration of these networks, examining whether the balance between within- and between-network connectivity was associated with BRS and statistically mediated the PGS–BRS association. We quantified this balance using the system–segregation index (SSI) in UKB (N = 27,729; Fig. 4d; see Supplementary Information for SSI distributions). SSI is calculated as the difference between within- and between-network connectivity, divided by within-network connectivity; a lower SSI value indicates greater relative integration.

Two sets of comparisons were used to calculate SSI for the CEN, SN, and DMN. We first compared connectivity within each network with its connectivity to the other two networks combined (one-to-triple SSI; Fig. 4e). The result showed that higher BRS was associated with lower SSI for the CEN (r = −0.047, P = 4.9 × 10⁻¹⁵) and DMN (r = −0.033, P = 3.5 × 10⁻⁸), indicating greater integration with the other two networks. The association for SN SSI was not significant (r = −0.009, P = 0.15; Fig. 4e, left). Among these three measures, only CEN SSI showed a significant indirect effect in the polygenic influence on BRS (standardized a × b = 0.63 × 10⁻³, P = 0.021; Fig. 4e, right). Higher PGS was associated with lower CEN segregation (a = −0.014, P = 0.021), and lower CEN segregation was associated with higher BRS (b = −0.046, P = 1.5 × 10⁻¹⁴). These findings suggest that CEN integration may help account for the PGS–BRS association, prompting us to examine its integration with the SN and DMN separately. In the pairwise analysis, SSI compared connectivity within each network with its connectivity to one other network (Fig. 4f). Five of six pairwise SSIs were associated with BRS (r = −0.019 to −0.044, all P ≤ 0.002; Fig. 4f, left). Only CEN integration with the SN showed a significant indirect effect (a × b = 0.57 × 10⁻³, P = 0.005; a = −0.017, P = 0.005; b = −0.035, P = 8.6 × 10⁻⁹; Fig. 4f, right). Notably, CEN integration with the DMN showed the strongest pairwise association with BRS (r = −0.044), but its indirect effect was not significant (P = 0.32). Together with the edge-level findings, these results support a potential mediating role for CEN integration, particularly with the SN, in the network-level pathway linking polygenic predisposition to resilience.

We next assessed whether these findings were consistent in GS. All seven SSIs associated with BRS in UKB showed the same direction in GS (N = 1,060), with DMN integration into the triple network reaching significance (r = −0.056, one-sided P = 0.034). In the mediation sample (N = 997), the indirect effects through CEN integration into the triple network and with the SN were numerically similar to those in UKB but did not reach significance, consistent with the limited power of the replication cohort (Fig. S8; Supplementary Information for details).

### Dissociable neural pathways underlying genetic and environmental contributions to resilience

The preceding analyses implicated CEN integration with the SN in the polygenic effect on resilience, while several other integration features were also associated with BRS. We next explored whether favorable environmental experiences might offset the lower resilience scores among individuals with lower PGS, and whether their network associations overlapped with or extended beyond the CEN–SN finding. We focused on childhood environmental quality (CEQ), given the relevance of early-life experiences to resilience in adulthood. Specifically, we examined CEQ–BRS associations across PGS levels and explored whether CEQ was linked to the CEN–SN integration measure or to additional network features associated with BRS. CEQ was indexed using reverse-scored items from the Childhood Trauma Screener in UKB and the Childhood Trauma Questionnaire–Short Form in GS, such that higher scores reflect a more supportive, non-maltreating early-life environment (Methods).

In UKB participants with CEQ data (N = 20,603), the PGS–CEQ association did not reach significance (β = 0.013, 95% CI −0.0003 to 0.027, P = 0.055; Fig. 5c). When modeled jointly, higher CEQ (β = 0.138, P = 4.4 × 10⁻^90^) and higher PGS (β = 0.066, P = 6.9 × 10⁻²²) were each associated with higher BRS. Their interaction was not significant (β = 0.003, P = 0.65; Fig. 5c). Mean BRS increased across PGS tertiles at each CEQ level, with approximately parallel gradients (Fig. 5b). Participants with both high PGS and high CEQ had the highest mean BRS (3.81), whereas those with both low PGS and low CEQ had the lowest (3.44). These findings are consistent with additive associations of PGS and CEQ with BRS, with no detected interaction in this model. The same pattern was observed in GS (N = 971), in which the PGS and CEQ were unrelated and additively associated with BRS (Fig. S9a; Supplementary Information for details).

**Fig. 5.**
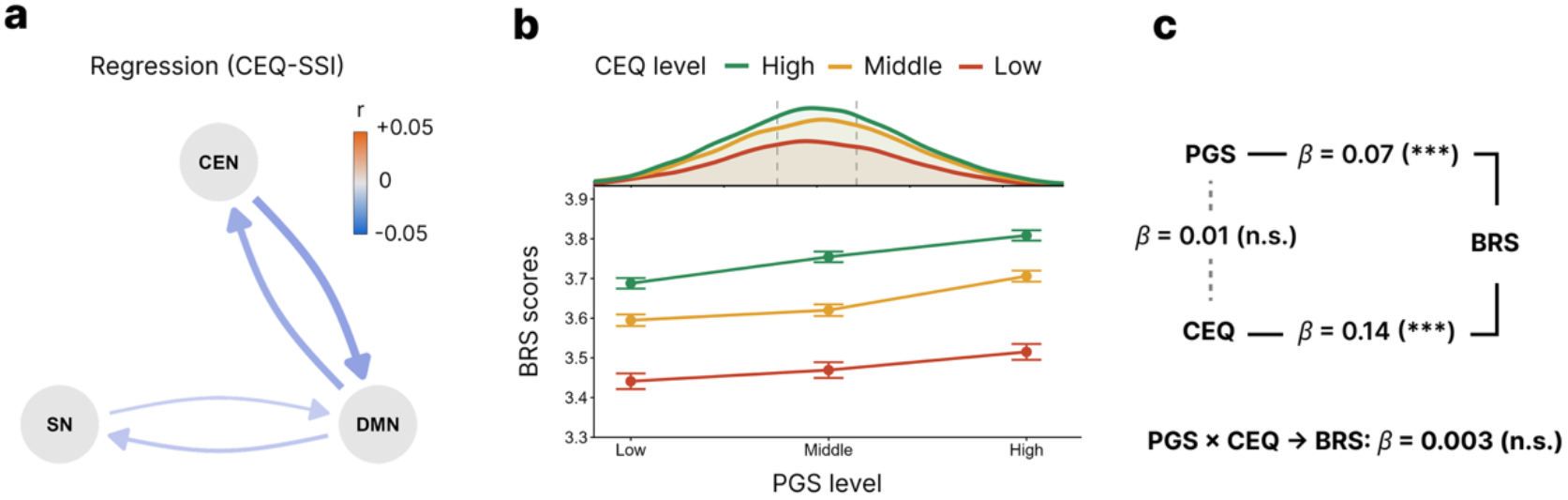
Associations of childhood environmental quality with triple-network integration and resilience. Analyses were performed in participants of the UKB imaging-genetics discovery cohort with complete CEQ data (N = 20,603). CEQ was calculated as the mean of the five Childhood Trauma Screener items after reverse-scoring the negatively worded items (range 1–5; higher values indicate a better childhood environment). a, Pairwise SSIs significantly associated with CEQ (P < 0.05); X → Y denotes the segregation of network X from Y. Arrow color and width indicate the partial correlation between CEQ and each rank-based inverse normal transformed SSI, adjusted for covariates. b, BRS scores (item mean, range 1–5; mean ± s.e.m.) across PGS tertiles, stratified by CEQ level. Top, PGS distributions for each CEQ group (kernel density scaled to the number of participants); dashed lines mark the PGS tertile cut points. c, Standardized regression coefficients (β) from regressing BRS on the PGS, CEQ and their interaction (solid lines), and from regressing CEQ on the PGS (dotted line). Models were adjusted for age, age squared and sex. ***P < 0.001; n.s., not significant.

We then examined whether CEQ was associated with the network integration features previously linked to BRS. Higher CEQ was associated with lower SSI for both networks in the CEN-DMN and SN-DMN pairs (four indices; *r* = −0.029 to −0.016, P = 2.6 × 10⁻⁵ to 0.025), indicating greater relative integration (Fig. 5a). In contrast, its association with CEN SSI relative to the SN (r = −0.013, P = 0.056) and SN SSI relative to the CEN (r = 0.002, P = 0.75) did not reach significance. Thus, CEQ was associated with all four resilience-related SSIs that did not carry the genetic effect, each in the direction of greater integration that accompanied higher resilience. These associations replicated in GS (Fig. S9b; Supplementary Information for details). These results suggest that genetic and early-environmental influences on resilience converge on triple-network integration through separable axes: a genetically shaped CEN–SN axis and an environmentally sensitive, DMN-centered axis.

## Discussion

Our findings portray self-reported resilience ability as both heritable and environmentally shaped, with each component linked to partly distinct integration signatures within the triple network. The BRS predicted symptom recovery after recent adversity, supporting its use as a population-scale measure of resilience. Its heritable component generalized across cohorts and was linked mainly to CEN–SN integration, whereas childhood environment related mainly to DMN-centered integration. Resilience may thus draw on a shared large-scale network through connections that differ by source of influence.

Many empirical studies have attempted to predict pre-traumatic mental health outcomes using resilience questionnaires^19–22^. However, a recent review concluded that there is still no evidence that resilience scales can predict mental health after adversity beyond baseline symptoms^10^. A common critique is that many studies lack (1) post-trauma baseline assessment^19–21^ and (2) a non-exposed comparison group^22,23^. A pre-trauma baseline ensures that the predictive value of resilience scale does not merely reflect its correlation with baseline mental health. A non-exposed group, in turn, confirms whether this predictive value is trauma-specific rather than a reflection of individual differences in mental health. In the present study, the BRS remained a significant predictor after accounting for both. Moreover, we showed that these adjustments matter: without them, the predictive value of the BRS was overestimated. These findings support the importance of controlling for both factors when validating resilience scales. This study has not fully resolved the phenotyping bottleneck in resilience research. Resilience is generally defined as better-than-expected mental health after adversity^2^, yet a post-trauma trajectory has at least two distinct components: the magnitude caused by the initial perturbation (reactivity/resistance) and the rate (bounce back) at which mental health returns to baseline. Here, we measure resilience as bouncing back, as this is the construct the BRS is designed to measure, whereas other studies have operationalized it as low trauma reactivity or resistance^10,24^. Both capture aspects of the post-traumatic recovery trajectory, but how each relates to RFs remains unclear. Due to the limitation of UKB data, the BRS was administered before recovery took place, so using it to predict bouncing back is temporally appropriate and rules out reverse causation. However, because it was administered after trauma exposure, BRS scores may already reflect individuals’ initial reactions to the trauma, which precludes a precise assessment of its relationship with trauma reactivity. Future studies that assess RFs before trauma exposure, together with repeated post-trauma assessments, would allow reactivity and bouncing back to be modeled as distinct outcomes and clarify how each relates to RFs.

Establishing that the BRS predicts recovery at population scale also made it possible to examine its neural correlates with adequate power. Previous rsfMRI studies of resilience included fewer than 300 participants, well below the thousands typically needed for reproducible brain–behaviour associations^9^. Testing the three main network hypotheses jointly in more than 27,000 participants with replication, we found that the triple network, which is thought to support flexible reallocation of attention and cognitive control^25^, showed the strongest positive association with the BRS. Notably, every tested edge within and between the CEN, SN and DMN was positively associated with the BRS, a pattern that previous studies would not have predicted. Resilience has most often been linked to stronger within-CEN, within-DMN and CEN–DMN connectivity, interpreted as better executive control over adversity and threat memories^6,26,27^. Findings involving the SN are mixed and appear to depend on the population studied. In adolescents, increases in within-SN connectivity mediated the association between childhood maltreatment and lower substance use^28,29^, and in non-clinical adults, stronger SN–DMN connectivity was associated with higher trait resilience in most studies^30–32^. By contrast, individuals with PTSD showed elevated connectivity both within the SN and between the SN and DMN^33,34^, interpreted as heightened threat detection and self-referential systems. Noting that these inconsistencies may stem from differences in the populations studied, and that our sample was drawn from the general population, we speculate that identifying less-resilient individuals by the presence of a mental disorder after trauma may reverse the apparent direction of associations, because such groups may reflect distinct brain states. Indeed, the proposed negative association between within-DMN connectivity and resilience is likewise a theoretical inference drawn from depression^15^. Evidence on SN– CEN connectivity is scarce. We propose that, at rest, stronger SN–CEN coupling may reflect a readiness to switch from detecting salient events to engaging executive control, a role attributed to the salience network in the triple-network model^25^. Such efficient flexible switching would support bouncing back after stress.

Our results link the polygenic architecture of resilience to large-scale brain organization. Heritability of the BRS was enriched in central nervous system tissue and in synaptic components, and the resulting polygenic score was associated with resilience partly through intrinsic connectivity of the triple network. Studying polygenic effects along the brain–phenotype axis has two implications. For causal inference, it addresses a limitation of conventional brain–phenotype association studies: because genetic variation is fixed at conception, the observed associations cannot arise from resilience or its correlates altering the genome. Genetic pleiotropy and genetically influenced environmental exposures nevertheless remain possible confounds, and future research is needed to test the robustness of this genetic pathway to resilience. In practice, localizing the genetic effect to a specific network axis, such as CEN–SN integration, can facilitate the development of mechanistically targeted interventions. In drug discovery, mechanisms with human genetic support are 2.6 times more likely to succeed in clinical development, and this advantage holds even for genetic associations of small effect^35^. By analogy, genetic evidence at the circuit level may help prioritize which brain correlates of resilience are worth targeting.

Taken together, our findings suggest that resilience is unlikely to rest on a single “resilience circuit”. Its neural correlates were distributed: all 45 triple-network connections and five of six pairwise integration indices were associated with the BRS, each with a small effect, and each of the three hypothesized systems explained unique variance in the BRS. One interpretation, which we offer as a hypothesis, is that resilience reflects a form of distributed robustness, in which many parts with distinct roles each contribute a small share to a function, so that a deficit in any single part has only a limited impact on the overall capacity^36^. Consistent with such partly independent contributions, polygenic predisposition was linked mainly to CEN–SN integration, whereas childhood environment was linked mainly to DMN-centred integration, and the two sources were nearly independent of each other. In such an architecture, an unfavorable influence on one route would leave other contributions intact, making the overall capacity less vulnerable to any single adverse factor, a property also attributed to degeneracy, in which structurally different elements yield the same output^37^. This interpretation should be weighed against two main limitations. First, most of the variation in resilience lies outside the connectivity we measured, and resting-state connectivity alone cannot establish that the brain achieves resilience through a distributed architecture. Future studies could combine multimodal measures, including structural, diffusion and task-based imaging as well as whole-brain rather than hypothesis-restricted networks, to test whether the distributed pattern holds across modalities and accounts for a larger share of resilience. Second, childhood environment was reported several years before the BRS was assessed, which reduces but does not eliminate concerns about reverse causation, because retrospective reports may still be confounded by individuals’ current appraisal of their past. More importantly, childhood environment is not randomly assigned and may be confounded by familial, socioeconomic and genetic factors that also shape adult resilience. A complementary strategy is to exploit natural experiments in which policy changes shift early-life environments independently of family characteristics. Applying such designs to resilience, and to the brain networks identified here, could isolate environmental contributions that are not confounded by familial or genetic factors.

By showing that a brief questionnaire captures recovery after real-life adversity, this study opens resilience to the sample sizes that genetic and neuroimaging research require. At this scale, heritable and early-life influences on resilience converged on the triple network, yet through partly distinct connections, consistent with a distributed rather than single-circuit architecture. Testing whether this distribution makes resilience robust, and whether its routes can be strengthened, will be essential for moving from population-level correlates to prevention.

## Methods

### Phenotypes and measures

**Resilience** was measured with the Brief Resilience Scale (BRS)^38^, a six-item questionnaire rated on a 5-point Likert scale. In the UK Biobank (UKB), BRS was collected in the second Mental Health Questionnaire^39^ (UKB field: 29175-29180; 2022-2023). In the Generation Scotland (GS), it was assessed during the STRADL remote follow-up and face-to-face assessment^40^ (2015-2019). An individual’s raw score is calculated as the average of the six items, three of which are reverse-scored. **Age** was defined as age at BRS completion (in UKB, age at recruitment [field 21022] plus the interval from the recruitment visit [field 53] to BRS completion [field 29208]) and **sex** as genetic sex (field 21001); both were provided by the study in GS. The following measures were available in UKB only unless stated otherwise. **Neuroticism** was the sum of 12 yes/no items assessed at recruitment (fields 1920–2030), set to missing if any item was missing. Participants reporting at least one of ten adverse events in MHQ2 (fields 29081–29090) “within the last 12 months” were classified as **trauma-exposed**; all others, including those reporting events only before that period, as unexposed. **Depressive and anxiety symptoms** over the preceding two weeks were measured at Time 0 (first assessment, 2006-2010) as the PHQ-4 (fields 2050-2080; 4 items, range 0–12), at Time 1 (MHQ2, 2022–2023) as the PHQ-9 (fields: 29002–29010) plus GAD-7 (field: 29058–29064) sum (range 0-48), and at Time 2 (online sleep questionnaire, 2022–2024) as the PHQ-4 (fields: 30484–30487, range 0–12). **Childhood environmental quality** (CEQ) was the mean of the five Childhood Trauma Screener items in MHQ1 (fields 20487–20491; 5-point Likert) and, in GS, of the 25 clinical items of its parent instrument, the Childhood Trauma Questionnaire–Short Form (CTQ-SF); negatively worded items were reverse-scored (UKB fields 20487, 20488, 20490), giving a range of 1–5 in both cohorts.

### Genotype quality control

In UKB, we use TOPMed-imputed data (field: 21007) as genotype, which includes 486,935 samples. In the quality control stage, we exclude samples with sex mismatches (field: 22019), and those identified as outliers for heterozygosity or missingness (field: 22027). Samples are subsequently restricted to the white ancestry subset, composed of individuals who self-identified as ‘British’, ‘Irish’, or ‘any other white background’ (field: 21000). In total, 446,293 participants are included in this subset. Here, we apply Bayesian Linear Mixed Model (BOLT-LMM) to conduct GWAS, which accounts for population stratification and hidden relatedness^41^, thus we do not adjust for detailed population structure or remove related individuals. Single Nucleotide Polymorphism (SNP) quality control was performed using PLINK (v1.9)^42^. We restrict the genetic variants to autosomal chromosomes, with a minor allele frequency (MAF) > 1%, a missingness rate < 2%, and conformity to Hardy–Weinberg equilibrium. Since the data were imputed, we included only variants with an imputation information score greater than 0.8. A total of 159,575 subjects passed genetic quality control and had BRS scores, age, sex, and genetic principal components (PC) records, constituting the GWAS discovery cohort in this study. Of these, 28,849 individuals had processed rsfMRI BOLD data, of whom 27,729 possessed complete parcellated matrices (detailed in the following section), forming the imaging-genetics discovery cohort. To derive polygenic scores (PGS) for this cohort without sample overlap, we re-ran the GWAS in the remaining 130,726 participants who had no processed rsfMRI data; we refer to this sample as the GWAS-for-PGS cohort.

In the GS cohort, 20,032 individuals had complete genotype data after quality control was performed by the GS:SFHS research group^43^. The original genotype data provided by GS is mapped to the hg19 reference genome. Because the TOPMed-imputed data is based on hg38, we used the liftover to update the GS coordinates to hg38 for proper alignment. Among them, a total of 8,702 had complete BRS scores and were used to conduct PGS validation.

### Functional connectivity of 3T resting-state fMRI

In UKB, resting-state fMRI data were processed in the official imaging pipeline^44,45^ and projected to regional time series using the Glasser cortical atlas^46^ (360 regions, 180 per hemisphere; field: 31016) together with the Tian Scale II subcortical atlas^47^ (32 regions, 16 per hemisphere; field: 31019). To test our a priori hypotheses regarding the neural substrates of mental resilience, we restricted downstream analyses to a predefined subset of these parcels. Cortical parcels were selected based on task-evoked functional responses followed by structural refinement, and subcortical parcels based on intrinsic functional organization (Supplementary Information). Table 1 lists the selected parcels and the anatomical ROIs to which they were assigned. For each pair of selected parcels, FC was computed as the Pearson correlation coefficient (r) between their BOLD time series. These r values were then Fisher r-to-z transformed to stabilize their variance and averaged within each anatomical ROI pair to yield anatomical-level FC. In the UKB, 27,729 participants had complete BRS scores, quality-controlled genotypes, and complete FC matrices; these formed the final analytic sample. For the Generation Scotland (GS) cohort, we obtained raw structural (T1-weighted) and resting-state fMRI data for 1,071 participants from the STRADL face-to-face assessment. Preprocessing of the T1-weighted and resting-state fMRI data followed the UKB pipeline as closely as possible, with adjustments for differences in MRI acquisition protocols (Supplementary Information). We applied the same parcellation atlases, ROI definitions, and FC computation. After preprocessing and parcellation, 1,060 participants had complete FC matrices and BRS scores; these participants were used to replicate the association between BRS and brain imaging. Of these, 997 also had genotype data from GS:SFHS and were used to replicate the imaging-genetics results.

**Table 1.**
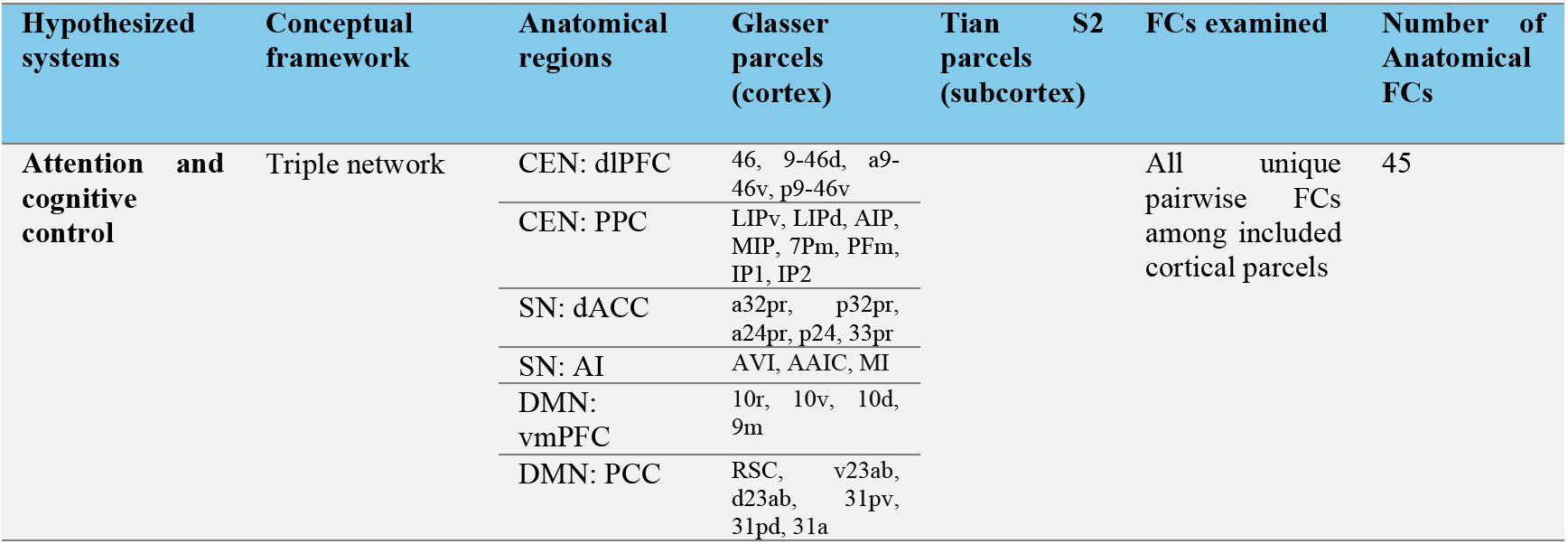
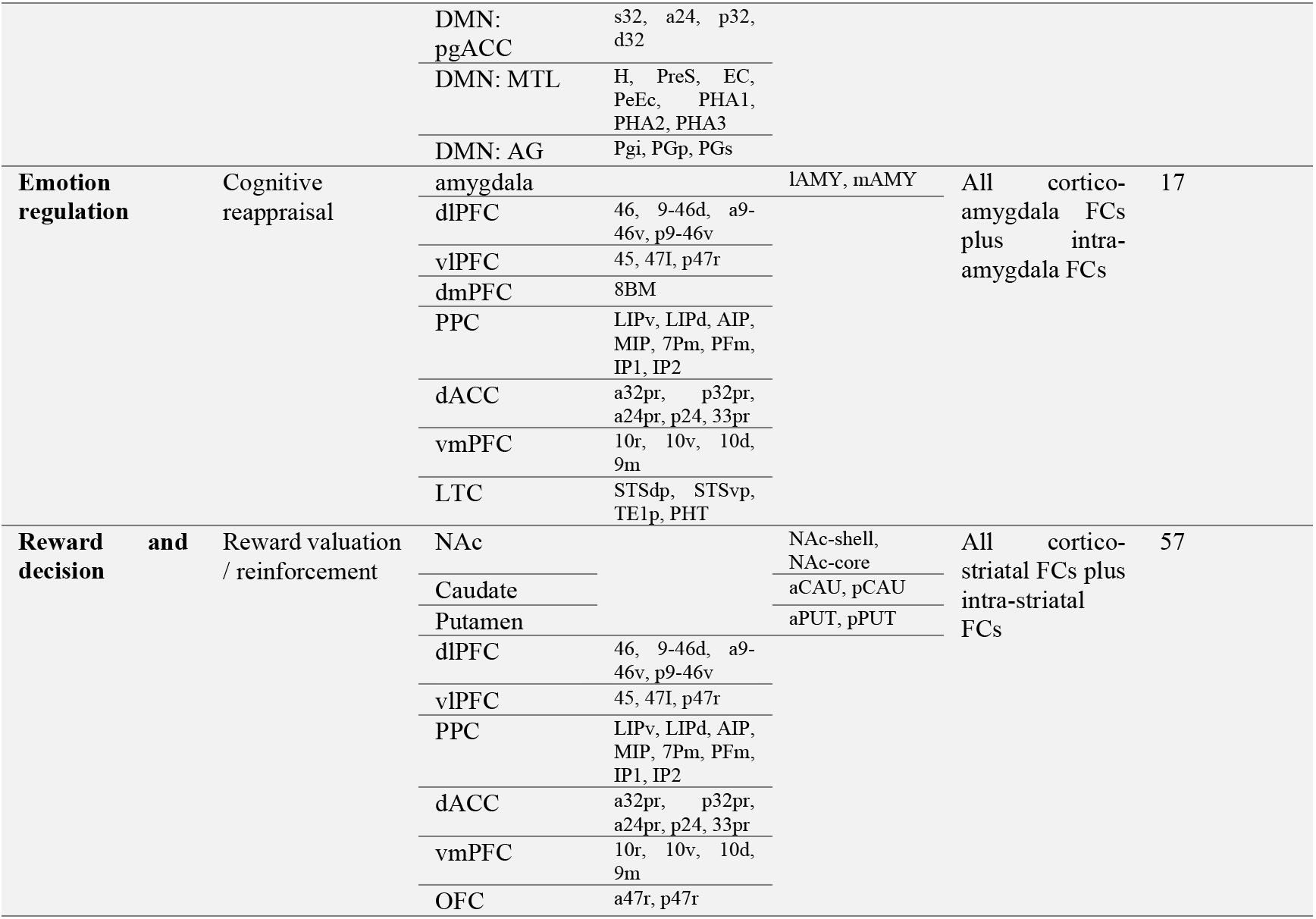
Target brain regions. Anatomical, atlas-based parcel mapping, and functional connectivity definitions for the predefined brain networks.

### BRS validation: trauma and symptom recovery

From the UKB GWAS discovery cohort (N = 159,575), we retained participants with a neuroticism score (N = 137,523) and complete data on distress at all three time points, BRS, the ten adverse-event items, age, sex and questionnaire completion dates (N = 112,622), and who completed Time 2 after Time 1, yielding an analytic sample of 110,346. BRS, neuroticism and distress at each time point were rank-based inverse normal transformed (INT) within this sample. In the main analysis, pooling all ten events, participants reporting at least one event within the past 12 months at Time 1 were classified as trauma-exposed (N = 8,319) and the remainder as controls (N = 102,027). The change in INT distress from Time 1 to Time 2 was regressed on BRS × trauma and neuroticism × trauma, adjusting for INT Time 0 and Time 1 distress, age, centered age squared and sex. To address confounding by pre-exposure distress, we ran two sensitivity analyses: (1) entropy balancing^48^, reweighting controls to match the exposed group’s mean Time 0 distress (exposed weight = 1); and (2) stratification by Time 0 distress (PHQ-4 0–2 normal, 3–5 mild, 6–12 moderate to severe), re-fitting the model within each stratum on the full-sample INT scale and testing heterogeneity with interaction × stratum terms in the full-sample model (Wald test, 2 d.f.). To examine how the BRS estimate depends on adjustment for Time 0 distress and on the comparison with controls, we also refitted the full-sample model without INT Time 0 distress, and estimated the BRS slope among exposed participants only, with and without INT Time 0 distress (all unweighted). All models used HC3 robust standard errors.

For illustration, we plotted raw distress trajectories for a single event, life-threatening injury or illness in the past 12 months, without restricting Time 0 distress (N = 105,359). Exposed participants and controls reporting none of the ten events were split at the median BRS (high ≥ 20 [raw score]; exposed high, n = 1,585; exposed low, n = 1,747), and Time 1 scores were rescaled to the PHQ-4 range (0–12). For each exposed group, changes from Time 0 to Times 1 and 2 are shown relative to BRS-matched controls. Both exposed groups were entropy-balanced to the Time 0 distress distribution of all exposed participants (one indicator per score from 1 to 8 and one for 9–12), and each control group was then balanced to the exposed group with the same BRS level on this distribution and on the mean and variance of BRS. Error bars are bootstrap standard errors (500 resamples, weights re-estimated in each). This figure is descriptive; inference relies on the regression models above.

### Genome-wide association study

We ran the GWAS using BOLT-LMM (v2.4.1)^49^ with BRS scores as phenotype. Two GWAS in this study were run separately, with identical settings, in the GWAS discovery cohort (n = 159,575) and in the GWAS-for-PGS cohort (n = 130,726). The model SNPs set used to derive genetic relationship matrix were generated by performing LD pruning on the quality-controlled genotypes with a window size of 100 SNPs, a step size of 5 SNPs, and an r² threshold of 0.1. The LD references are based on European-ancestry samples from 1000 Genomes Project phase 3. When conducting the GWAS, common covariates include age, centered age squared, sex, the first 20 genetic principal components (field: 22009), and genotype batch (field: 22000) are all considered.

### Polygenic scores

PGS were computed with LDpred2^50^ using the precomputed HapMap3+ LD reference^51^, merged with GWAS summary statistics on hg38 positions. In the UKB imaging-genetics discovery cohort (n = 27,729), PGS were derived from the GWAS-for-PGS summary statistics (n = 130,726). In GS, two sets were derived: from the GWAS discovery cohort (n = 159,575), to validate the primary GWAS signal in an independent sample, and from the GWAS-for-PGS cohort, to replicate the UKB imaging-genetics analyses. We applied both the infinitesimal model (LDpred2-inf; all variants causal, p = 1) and LDpred2-auto, which infers p and h² by MCMC, running 30 Gibbs chains initialized with p log-spaced from 10⁻⁴ to 0.2. h² was fixed (inf) or initialized (auto) at the LDSC estimate. Predictive performance was assessed by regressing BRS on each PGS in both cohorts, adjusting for age, centered age squared and sex.

### SNP-level heritability and tissue-specific enrichment

Following the GWAS results, LD score regression (LDSC)^52^ is used to estimate the SNP-based heritability of resilience for standard SNPs mapped to HapMap3, with European 1000 Genomes Project phase 3 as LD reference panel. Functional heritability partitioning is subsequently performed to examine functional and tissue-specific enrichments^53^. The latest version of Baseline-LD model (v2.2) is incorporated as base model, which comprises 97 overlapping functional and LD-related annotations^54^. Building upon this baseline model, we further perform cell-type-specific enrichment analyses to identify the biological tissues driving the heritability. Ten distinct cell-type groups, defined by specific histone modifications, were evaluated by adding each group’s annotation individually to the full baseline-LD model^53^. The significance of each cell-type group’s specific contribution is assessed using the regression coefficient P-values. P-values were corrected for multiple testing using the Benjamini–Hochberg false discovery rate (FDR) within each dataset.

### Gene-level over-representation analysis and cell type analysis

Gene-level association analysis was performed using MAGMA (v1.10)^55^ using the SNP-wise mean model. SNPs were mapped to genes using the NCBI build 38 gene locations provided with MAGMA, with gene windows extended 35 kb upstream and 10 kb downstream of gene boundaries, and linkage disequilibrium was estimated from the 1000 Genomes Phase 3 European reference panel. Subsequently, we perform an Over-Representation Analysis (ORA) by using *clusterProfiler* (v4.16.0)^56^, using a one-sided hypergeometric test on 2,038 MAGMA-identified significant candidate genes (e.g., P<0.05) across the Cellular Component (CC) category of the Gene Ontology (GO)^57^, as provided by MSigDB (v2025.1.Hs), and the 18,828 genes with MAGMA results served as the background universe. Of the 1,042 GO-CC gene sets, 642 contained 10–2,000 background genes and at least one input gene and were tested; P-values were FDR corrected.

Cell-type specificity was tested with the FUMA Cell Type module^58^, using the MAGMA gene-level results above as input so that gene annotation and windows were consistent. This tests whether genes more strongly associated with BRS are more specifically expressed in each cell type, conditioning on average expression across all cell types in the dataset. We used single-cell RNA-seq data from the Linnarsson mouse nervous system atlas^59^ (265 cell types) and the Allen human brain atlas^60^ (75 middle temporal gyrus [MTG] and 12 lateral geniculate nucleus [LGN] cell types). In the Linnarsson mouse nervous system atlas, cell types were grouped by their principal neurotransmitter (glutamatergic, GABAergic, dopaminergic, serotonergic, cholinergic, or other) and by anatomical/developmental origin (telencephalon, diencephalon, mesencephalon, cerebellum, hindbrain, spinal cord, peripheral nervous system, or non-regional glia/vascular cells), following the cluster nomenclature of the publication^61^. In the Allen human brain atlas, following the taxonomy of the publication^60^, human cell types were grouped by region (MTG or LGN) and by major cell class: glutamatergic, GABAergic, and non-neuronal cells, which were further divided into astrocytes, microglia, oligodendrocyte-lineage cells (oligodendrocytes and OPCs), and endothelial cells. P-values were FDR corrected in Linnarsson. Because no human cell type survived FDR correction, cell types with nominal P < 0.05 are highlighted as suggestive associations.

### Brain edge-level of functional connectivity correlation

119 anatomical-level FCs among three hypothesized systems (attention/cognitive control, 45 FCs; emotion regulation, 17 FCs; reward/decision-making, 57 FCs) associated with mental resilience, were separately fitted by linear regression on INT BRS scores (N = 27,729) in UKB imaging-genetics discovery cohort. Each FC feature was defined as the mean of parcel-level FC between two anatomical ROIs. Each model regressed resilience onto a single FC, controlling for age, mean-centered age squared, sex, and two rsfMRI data-quality measures: mean absolute head motion and temporal signal-to-noise ratio (tSNR). Head motion was indexed by the UK Biobank imaging-derived phenotype for mean absolute head motion (UKB field: 24438), and tSNR by the inverse tSNR of the artefact-cleaned preprocessed BOLD data (UKB field: 25744), which we converted to tSNR. Because both measures were right-skewed, they were log-transformed (log head motion; log tSNR = −log[inverse tSNR]) and entered as continuous covariates. Partial correlation coefficients were derived from the t statistics, and P-values were corrected using FDR. Because the three networks comprise different numbers of FCs, the number of significant edges is not directly comparable across networks. We therefore evaluated each network at the network level using nested model comparisons with partial F-tests, which test a set of FCs jointly and account for the number of FCs through the numerator degrees of freedom. The full model regressed resilience onto all 119 FCs simultaneously, together with the same covariates (age, centered age squared, sex, log head motion, and log tSNR). For each target network, the full model was compared with a reduced model from which only that network’s FCs were removed. The partial F statistic, F = [(RSS_reduced − RSS_full)/q] / [RSS_full/df_full], where q is the number of FCs in the target network, tests whether the target network explains variance in resilience beyond the covariates and the other two networks. P-values were Bonferroni corrected.

Replications were tested in imaging replication cohort in GS (N = 1,060). For cross-cohort comparison, FCs were defined using the same anatomical ROIs (119 FCs). We performed edge-wise regression with the same model as in UKB, regressing INT BRS scores onto each FC while controlling for age, age squared, sex, log head motion, log tSNR, and scanning site (because GS participants were scanned at two sites, Aberdeen and Dundee, using different scanners and acquisition parameters). Partial correlation coefficients were derived as in UKB, and edge-wise partial correlations were compared between the two cohorts. To assess whether effects replicated in the direction observed in UKB, we computed, for each network, the mean of the GS edge-wise partial correlations after aligning each FC to the sign of its UKB estimate (i.e., the mean of sign[r_UKB] × r_GS), such that positive values indicate replication in the UKB direction. Because FCs within a network are correlated, significance was assessed by permutation: INT BRS scores were randomly permuted 10,000 times (within each imaging site), edge-wise partial correlations were re-estimated with the same covariates in each permutation, and the sign-aligned mean was recomputed. The one-sided permutation P-value was computed as (1 + the number of permutations with a sign-aligned mean at least as large as observed) / (1 + 10,000). For each network, we also counted the number of FCs whose GS partial correlation had the same sign as in UKB. This count was tested with a one-sided exact binomial test (match probability = 0.5 per FC), which assumes independent FCs, and with the same within-site permutation procedure.

### Brain multivariate functional connectivity modeling

Network-specific predictive models in UKB imaging-genetics discovery cohort (N = 27,729) were trained, and the resulting weights were applied, without refitting, to GS (N = 1,060). For each network, INT BRS scores and each FC were first residualized on the covariates (age, mean-centered age squared, sex, log head motion, and log tSNR) and standardized. Three models were trained: (1) connectome-based predictive modelling (CPM), in which FCs whose correlation with residualized resilience reached P < 0.01 were selected and combined into a network-strength score by summing positively associated FCs and subtracting negatively associated FCs; (2) partial least squares regression with a single latent component (PLS); and (3) ridge regression, with the penalty parameter selected by 10-fold cross-validation (minimum mean squared error). In GS, the 119 FCs were computed using the same ROI definitions as in the UKB main analysis, residualized on the GS covariates (age, age squared, sex, imaging site, log head motion, and log tSNR), and standardized; a predicted score was then obtained as the weighted sum of FCs using the UKB-derived weights. Predictive performance was quantified as the partial correlation between the predicted score and INT BRS, adjusting for the same covariates. One-tailed P-values are reported (direction of association was specified by the UKB-trained models). To compare models, we computed Pearson correlations between the GS predicted scores of different models and compared their FC weights after scaling each model’s weights by its maximum absolute weight within each network.

### Functional connectivity mediation of polygenic effects

Edge-wise mediation was tested in UKB imaging-genetics discovery cohort (N = 27,729) with the PGS as the predictor (X), each of the 119 FCs as the mediator (M), and INT BRS as the outcome (Y), adjusting for the same covariates as the edge-wise regression. The a path was estimated by regressing the FC on the PGS, the b path by regressing INT BRS on the FC and the PGS, and the total effect (c) by regressing INT BRS on the PGS. The indirect effect was quantified as a × b and, for comparison across FCs and cohorts, as the completely standardized indirect effect (a × b × SD_X / SD_Y). Because a × b is not normally distributed, significance was assessed with the Monte Carlo method, which performs comparably to bootstrapping in large samples at a much lower computational cost^61^: a and b were drawn 1,000,000 times from normal distributions defined by their estimates and standard errors, the 95% confidence interval was taken from the 2.5th and 97.5th percentiles of their products, and the two-sided P-value as twice the smaller tail proportion on either side of zero (with a +1 correction). P-values were FDR corrected within each network.

Mediation was replicated in imaging-genetics replication cohort (N = 997) using the same FCs, models, and Monte Carlo procedure, with scanning site as an additional covariate, and the standardized indirect effect of each FC was compared between cohorts. To test whether the direction and magnitude of the indirect effects replicated, we computed, for each network, the number of FCs whose GS indirect effect had the same sign as in UKB and the mean GS standardized indirect effect after aligning each FC to the sign of its UKB estimate; significance was assessed by permuting the PGS within imaging site (10,000 permutations) and re-estimating all indirect effects (one-sided P = [1 + number of permutations ≥ observed] / [1 + 10,000]).

To estimate the proportion of the PGS–resilience association jointly mediated by the attention/cognitive control network (H1), we fitted a parallel multiple mediator model in UKB imaging-genetics discovery cohort (N = 27,729) using the 13 H1 FCs with significant edge-wise indirect effects (FDR < 0.05): each a_i path was estimated separately, and all b_i paths simultaneously by regressing INT BRS on the 13 FCs and the PGS, with the same covariates as above. We estimated the total indirect effect (Σ a_i × b_i) and the total proportion mediated (PM = Σ a_i × b_i / c). 95% confidence intervals were obtained by percentile bootstrapping (5,000 resamples of participants).

### Network integration/segregating analysis from functional connectivities

To characterize the balance between within- and between-network connectivity of the triple network (CEN, SN and DMN), we computed a segregation–integration index (SSI) from the 45 H1 FCs in the UKB imaging-genetics discovery cohort (N = 27,729), the GS imaging replication cohort (N = 1,060) and the GS imaging-genetics replication cohort (N = 997). The FCs were grouped into three within-network blocks (CEN, 3 FCs; SN, 3 FCs; DMN, 15 FCs) and three between-network blocks (CEN–SN, 4 FCs; CEN–DMN, 10 FCs; SN–DMN, 10 FCs). Following the system segregation framework, negative FCs were set to zero, and each block was summarized as the mean of its FCs. The pairwise SSI of network X relative to network Y was defined as (FC_X − FC_X–Y) / FC_X, where FC_X is the within-network FC of X and FC_X–Y is the between-network FC of X and Y. The one-to-triple SSI of network X was defined as (FC_X − FC_X–other) / FC_X, where FC_X–other is the average of the between-network FCs of X with the other two networks, weighting the two networks equally. Higher SSI indicates greater segregation, and lower SSI greater integration, of a network with the others. This yielded nine SSIs (three one-to-triple and six pairwise). Each SSI was rank-based inverse normal transformed within each cohort before analysis (because SSI is a ratio that might become unstable when within-network FC approaches zero).

The association of each SSI with resilience was tested by linear regression of INT BRS scores on the SSI, using the same covariates as the edge-wise regression (age, mean-centered age squared, sex, log head motion and log tSNR) in UKB (N = 27,729). Replication was tested in the GS imaging replication cohort (N = 1,060) with the same models, with scanning site as an additional covariate. For SSIs significantly associated with BRS in UKB, one-sided P-values were computed in the direction observed in UKB. Mediation through SSI was tested with the same procedure as the edge-wise mediation analysis. In UKB (N = 27,729), the PGS was the predictor (X), each SSI the mediator (M), and INT BRS the outcome (Y), with the same covariates. The analysis was repeated in the GS imaging-genetics replication cohort (N = 997) with scanning site as an additional covariate, and the standardized indirect effects were compared between cohorts.

### Association between childhood experiences, resilience, and brain systems

Analysis was conducted on UKB imaging-genetics discovery cohort and the GS imaging-genetics replication cohort with available CEQ data. Of the 27,729 UKB participants, 20,603 had complete CEQ data; of the 997 GS participants, 971 had complete CEQ data.

To compare the contributions of the PGS and CEQ to resilience, BRS scores were regressed on the PGS, CEQ and their interaction (PGS × CEQ), adjusting for age, age squared and sex, with imaging site as an additional covariate in GS. The association between the PGS and CEQ was tested by regressing CEQ on the PGS with the same covariates. The PGS, CEQ and BRS scores were standardized, so that coefficients are standardized β. For visualization, UKB participants were grouped by PGS tertile and by CEQ level.

The association of CEQ with triple-network segregation was tested with the nine SSIs defined above, rank-based inverse normal transformed within each cohort. Each SSI was regressed on CEQ with the same covariates as the edge-wise regression (age, mean-centered age squared, sex, log head motion and log tSNR), with scanning site as an additional covariate in GS. Partial correlation coefficients and their 95% confidence intervals were derived as in the SSI–resilience analysis. In GS, participants whose within-network FC was zero after negative FCs were set to zero were excluded from the SSIs of that network (N = 971). For SSIs significantly associated with CEQ in UKB, one-sided P-values were computed in the direction observed in UKB.

## Author contributions

C.-Y.H., Y.-Y.Y., T.-Y.W., H.-C.Y., and F.-H.L. conceived the study. C.-Y.H., Y.-Y.Y. and T.-Y.W. designed the study. T.-Y.W. performed the analyses and wrote the first draft of the manuscript. Y.-Y.Y. and Y.-J.L. reviewed the analyses and provided feedback throughout the study. C.-Y.H., H.-C.Y., and F.-H.L. supervised the study. Y.-Y.Y. and C.-Y.H. revised the manuscript. C.-Y.H. acquired funding.

## Acknowledgement

We are grateful to all the participants who took part in the UK Biobank and Generation Scotland, and the whole team of UK Biobank and Generation Scotland, which includes interviewers, computer and laboratory technicians, clerical workers, research scientists, volunteers, managers, receptionists and nurses. This research has been conducted using the UK Biobank Resource under Application Number 465546 and Generation Scotland data under GS Application Number GS25578. Generation Scotland received core support from the Chief Scientist Office of the Scottish Government Health Directorates [CZD/16/6] and the Scottish Funding Council [HR03006] and is currently supported by the Wellcome Trust [216767/Z/19/Z]. Genotyping of the GS:SFHS samples was carried out by the Genetics Core Laboratory at the Wellcome Trust Clinical Research Facility, Edinburgh, Scotland and was funded by the Medical Research Council UK and the Wellcome Trust (Wellcome Trust Strategic Award “STratifying Resilience and Depression Longitudinally” (STRADL) Reference 104036/Z/14/Z).

This work was funded by the Academia Sinica Scholar Award (ASSA), Academia Sinica, Taiwan (AS-ASSA-113-04). We thank Chien-Yu Lai and Sheng-Kai Chang for helpful comments on causal inference and statistical testing, and Yen-Tsung Huang for advice on the interpretation of the mediation analyses.

## Supplementary Information

This document provides supplementary notes, figures, and references supporting the main text. **Table S1 lists each supplementary figure together with its section** in this document and the corresponding sections of the main-text Results and Methods.

**Table S1.** Guide to the Supplementary Information. Each supplementary figure (Item) is listed with the title of its section in this document (Context) and the related sections of the main-text Results and Methods. A dash indicates that the figure has no corresponding Results section.

| **Item** | **Context** | Related section: Results | Related section: Methods |
|---|---|---|---|
| **Fig. S1** | **Cohort overview** | Cohort overview and phenotypic characterization of BRS | •Genotype quality control<br>• Functional connectivity of 3T resting-state fMRI<br>•BRS validation: trauma and symptom recovery |
| **Fig. S2** | **Sensitivity analysis of BRS validation** | BRS predicts resilient outcome | BRS validation: trauma and symptom recovery |
| **Fig. S3** | **Polygenic scores validation** | Genetic factors and cell types through genome sequencing data | Polygenic scores |
| **Fig. S4** | **Genetic enrichment analysis on human brain** | Genetic factors and cell types through genome sequencing data | Gene-level over representation analysis and cell type analysis |
| **Fig. S5** | **Multivariate rsfMRI predictive model** | Brain networks assessed by resting-state fMRI | Brain multivariate functional connectivity modeling |
| **Fig. S6** | **Replication of edge-wise association** | Brain networks assessed by resting-state fMRI | Brain edge-level of functional connectivity correlation |
| **Fig. S7** | **Replication of edge-wise mediation** | Brain network mediation of genetic resilience factors | Mediation effects of functional connectivities to resilience polygenicity |
| **Fig. S8** | **Replication of system integration** | Relative balance between brain networks as resilience factors | Networks integration/segregating analysis from functional connectivities |
| **Fig. S9** | **Replication of childhood environmental effect** | Dissociable neural pathways underlying genetic and environmental contributions to resilience | Association between childhood experiences, resilience, and brain systems |
| **Fig. S10** | **MRI data processing in GS** | - | Functional connectivity of 3T resting-state fMRI. |
| **Fig. S11** | **Brain hypothesis** | Brain networks assessed by resting-state fMRI | Functional connectivity of 3T resting-state fMRI. |

### Cohort overview

Fig. S1 maps the participants included in each analysis onto the data-collection timeline of each cohort. Within UKB, the PGS was derived from a GWAS restricted to participants without rsfMRI data, so that the imaging-genetics discovery cohort does not overlap with the GWAS used to construct the PGS; the BRS validation cohort is a subset of the GWAS discovery cohort used for behavioral analyses only. GS provides an independent replication sample at each level of analysis: PGS validation, BRS–FC associations and imaging-genetics associations.

**Fig. S1.**
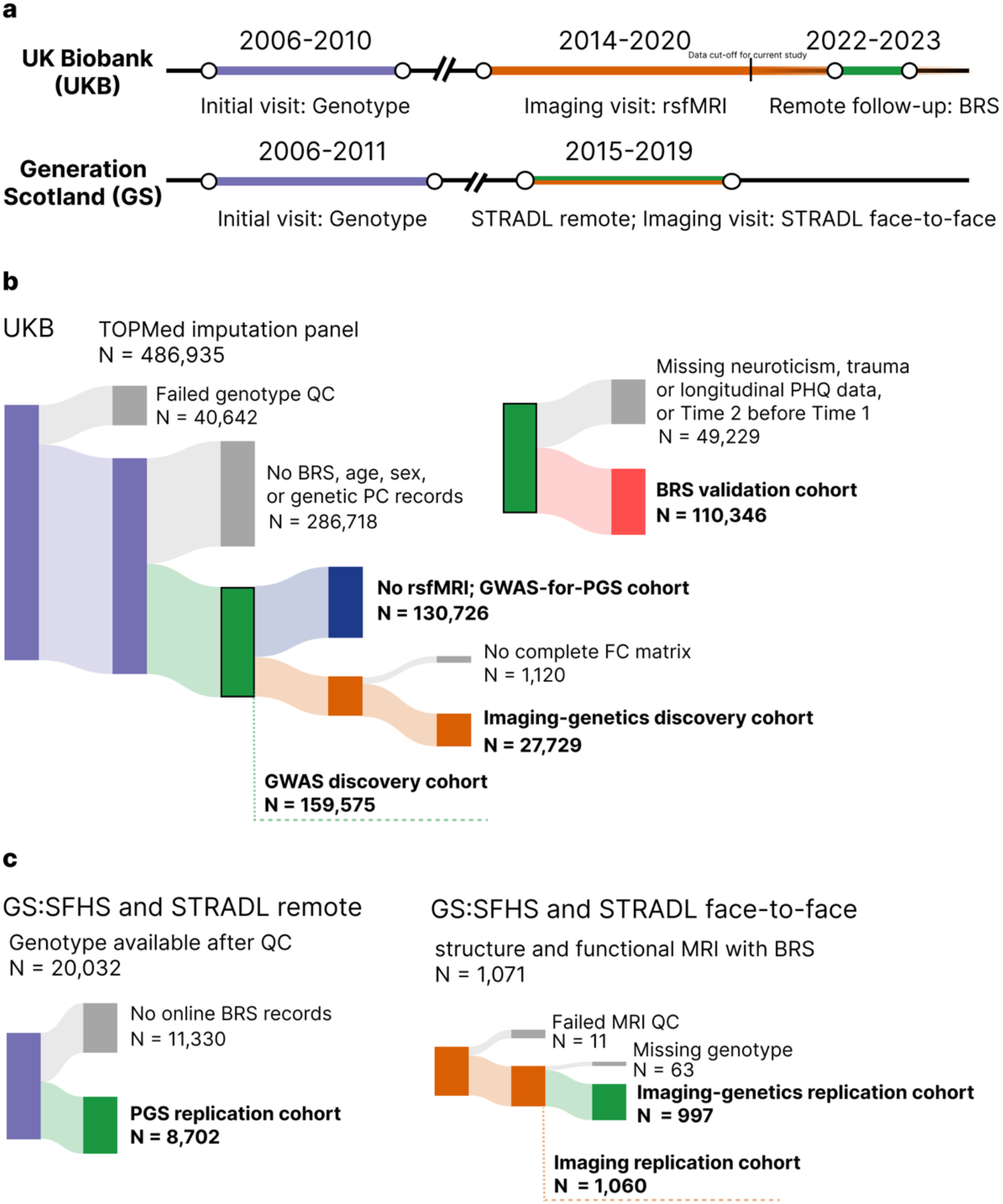
Study timeline and sample selection. **a**, Timeline of data collection in UK Biobank (UKB) and Generation Scotland (GS). In UKB, genotypes were obtained at the initial visit (2006–2010), resting-state functional MRI (rsfMRI) at the imaging visit (2014–2020), and BRS at remote follow-up (2022–2023). In GS, genotypes were obtained at the initial visit (2006–2011), and BRS and MRI through the STRADL remote and face-to-face assessments (2015–2019). **b**, Sample selection in UKB. Of 486,935 participants with TOPMed-imputed genotypes, 159,575 passed genotype quality control (QC) and had BRS, age, sex and genetic principal component (PC) data (GWAS discovery cohort). This cohort was split into participants without rsfMRI, used for GWAS to derive the polygenic score (PGS; N = 130,726), and participants with complete functional connectivity (FC) matrices (imaging-genetics discovery cohort; N = 27,729). Participants from the GWAS discovery cohort with complete neuroticism, trauma and longitudinal PHQ data, and with Time 2 completed after Time 1, formed the BRS validation cohort (N = 110,346). **c**, Sample selection in GS. Of 20,032 genotyped GS: SFHS participants, 8,702 with BRS from STRADL remote formed the PGS replication cohort. Of 1,071 STRADL face-to-face participants with structural and functional MRI and BRS, 1,060 passed MRI QC (imaging replication cohort), of whom those with genotype data formed the imaging-genetics replication cohort.

Note that BRS was collected several years after imaging in UKB but within the same assessment period as imaging in GS (Fig. S1a). Exclusion criteria are described in Methods (Genotype quality control; Functional connectivity of 3T resting-state fMRI; BRS validation), and MRI processing in GS in Supplementary Section of “MRI data processing in GS” (Fig. S10).

### Sensitivity analysis of BRS validation

#### Entropy balancing on pre-exposure distress (Fig. S2a, b)

Trauma-exposed participants differed from unexposed participants before exposure. Their mean Time 0 PHQ was higher (1.85 vs 0.95; SMD = 0.46), they were less often symptom-free (SMD for Time 0 PHQ = 0 = −0.39), they had higher neuroticism (SMD = 0.21), and they had lower BRS (SMD = −0.23) (Fig. S2a). The main model adjusts for Time 0 PHQ linearly. When groups differ this much, however, the interaction estimate can depend on whether that linear adjustment is correctly specified across the whole range of baseline distress. We therefore reweighted unexposed participants with entropy balancing^1^, so that the comparison no longer relies on the regression model alone to correct for the baseline difference. For unexposed participants with Time 0 PHQ X_i_, entropy balancing chooses weights w_i_ that stay as close as possible to uniform weights q_i_ = 1/n₀ while exactly matching the exposed group’s mean X̄₁:

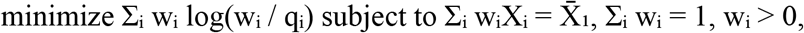

where sums run over the n₀ unexposed participants. The solution has the closed form

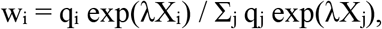

where λ is found by solving the convex dual problem. Exposed participants kept a weight of 1, so the estimand refers to the exposed population (average treatment effect on the treated). Unlike propensity-score weighting, entropy balancing does not require a model of exposure, and it achieves exact balance on the chosen moment rather than balance that has to be checked afterwards.

After weighting, mean Time 0 PHQ was identical in the two groups (SMD = 0.00). The weights were not targeted at any other covariate, yet all remaining differences fell within |SMD| < 0.1: neuroticism 0.04, age 0.06, sex −0.09, and BRS −0.05 (Fig. S2a). The one exception was a small residual difference in the share of participants without symptoms (SMD = −0.11), and the squared term was close behind (SMD = −0.08). The weights were moderately dispersed: the effective sample size of the unexposed group was 81,576 of 102,027 (80%). Weighting shifted the unexposed reference toward participants with higher baseline distress. The BRS slope in this reference group was correspondingly steeper (β = −0.075 vs −0.067 unweighted), whereas the slope among exposed participants was essentially unchanged (β = −0.102 vs −0.104; Fig. S2b). The BRS × trauma interaction was therefore attenuated (β = −0.027, 95% CI [−0.043, −0.011], vs −0.037). The attenuation reflects a different reference group, not a change among exposed participants. The interaction remained significant (P = 8.9 × 10⁻⁴). The neuroticism × trauma interaction remained null (β = −0.003, P = 0.73).

**Fig. S2.**
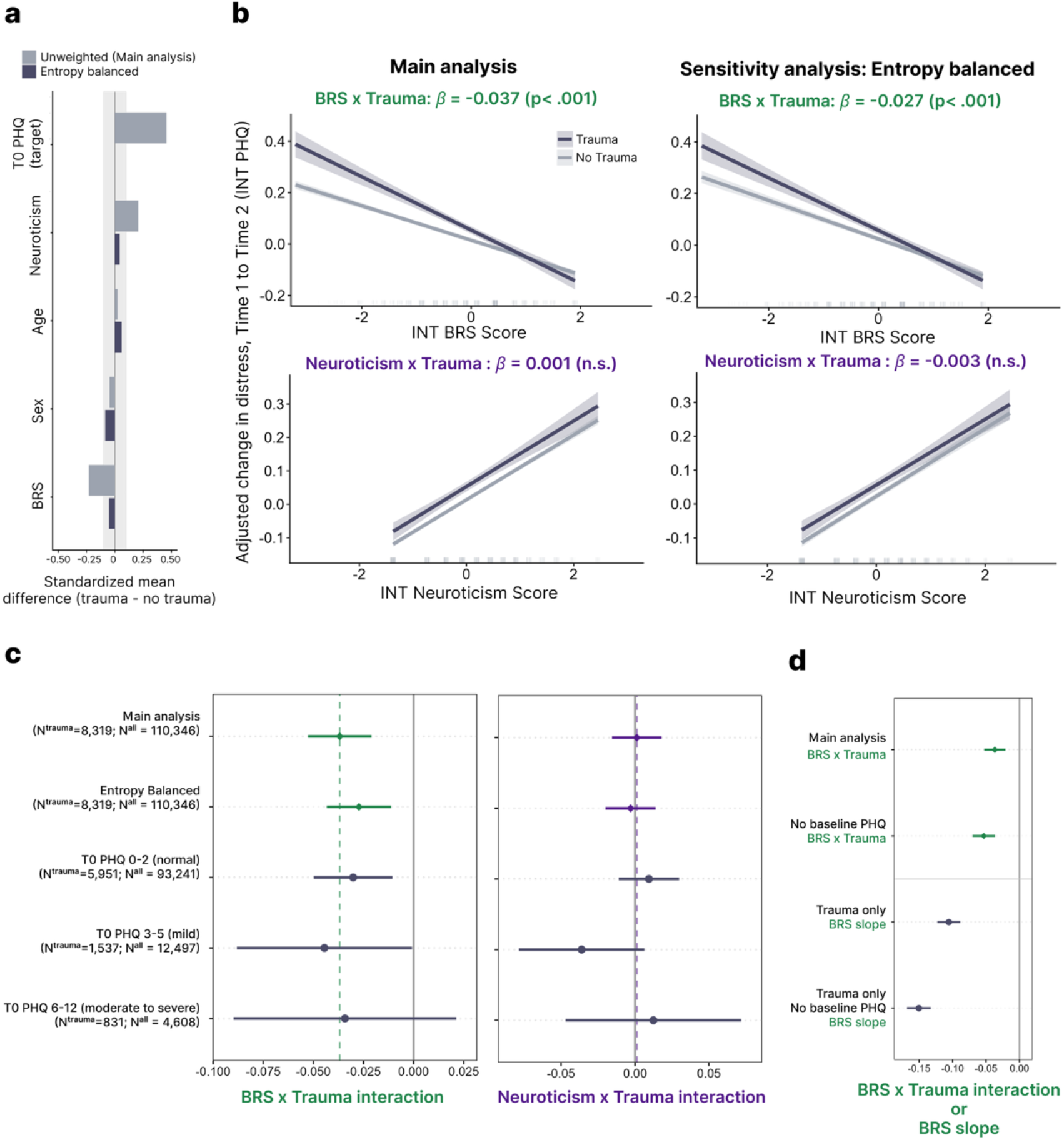
Sensitivity analyses of the BRS × trauma interaction on symptom recovery. **a**, Covariate balance between trauma-exposed (n = 8,319) and unexposed (N = 102,027) participants before (unweighted; main analysis) and after entropy balancing on Time 0 PHQ, shown as standardized mean differences (SMD; trauma − no trauma). The shaded band indicates |SMD| < 0.1. **b**, Predicted change in psychological distress from Time 1 to Time 2 (INT PHQ) as a function of inverse-normal transformed (INT) BRS (top) and neuroticism (bottom) scores, stratified by trauma exposure, in the main analysis (left; replicated from Fig. 1g for comparison) and after entropy balancing (right) (N = 110,346). Models adjusted for Time 1 and Time 0 PHQ, sex, age and centered age squared; continuous covariates were held at their sample means and sex at its reference category (male). Coefficients are on the INT scale. **c**, BRS × trauma (left) and neuroticism × trauma (right) interaction estimates in the full sample (main analysis and entropy balanced) and within Time 0 PHQ severity strata (PHQ-4: 0–2 normal, 3–5 mild, 6–12 moderate to severe; the moderate and severe bands were merged because of small numbers). Stratum estimates are unweighted. Dashed vertical lines indicate the main-analysis estimates. Heterogeneity across strata: BRS × trauma × stratum F(2) = 0.28, P = 0.76; neuroticism × trauma × stratum F(2) = 2.00, P = 0.14. **d**, BRS estimates under simpler model specifications (unweighted): the BRS × trauma interaction in the main model and without adjustment for Time 0 PHQ (N = 110,346), and the BRS slope among trauma-exposed participants only, with and without adjustment for Time 0 PHQ (n = 8,319). The trauma-only slope also includes the association between BRS and symptom change shared with unexposed participants. Shaded bands in b and error bars in c and d represent 95% confidence intervals based on heteroskedasticity-consistent (HC3) standard errors.

#### Stratification by pre-exposure distress (Fig. S2c)

As a second, model-free check, we fitted the main model separately within Time 0 PHQ-4 severity bands (0–2 normal, 3–5 mild, 6–12 moderate to severe^2^). Within a band, exposed and unexposed participants are compared at similar levels of baseline distress. Unlike a single linear adjustment, stratification also lets the interaction itself differ by baseline severity. This addresses the concern that the interaction is driven by participants with high baseline distress, for example through regression to the mean or ceiling effects. Heterogeneity was tested by adding three-way interaction terms to the full-sample model.

The BRS × trauma interaction was similar in all groups (β = −0.030, −0.045 and −0.034; heterogeneity F(2) = 0.28, P = 0.76). The main limitation of this approach is reduced precision. Stratification splits the sample unevenly: 93,241 participants were in the lowest band, but only 4,608 (831 exposed) were in the highest. The heterogeneity test therefore has limited power to detect modest differences across strata.

#### Consequences of incomplete adjustment (Fig. S2d)

Many studies of resilience scales do not include an unexposed comparison group, and many do not adjust for pre-exposure mental health. Reviews in the field have called for both^3,4^. To quantify what these choices imply, we re-estimated the BRS effect under simpler specifications. With an unexposed group, the estimand remains the BRS × trauma interaction: the difference in BRS slope between exposed and unexposed participants. Without an unexposed group, exposure does not vary, so the interaction cannot be estimated. The only available estimate is then the BRS slope among exposed participants, which is how most studies of resilience after adversity analyse their data. This slope corresponds to the sum of the BRS main effect and the BRS × trauma interaction in the full model. Indeed, the slope implied for exposed participants by the full model (−0.104) closely matched the slope estimated in exposed participants alone (−0.106).

Both simplifications inflated the estimate. Omitting Time 0 PHQ increased the interaction by 45% (β = −0.053 vs −0.037). Higher BRS is associated with lower pre-exposure distress, so without this adjustment part of the baseline difference is attributed to BRS. Restricting the analysis to exposed participants gave a slope about three times larger than the interaction (β = −0.106). About two-thirds of this slope (β = −0.067) was also present in unexposed participants, so it reflects the general association between BRS and symptom change, not recovery from the event. Combining both simplifications gave a slope about four times larger than the interaction (β = −0.150). In other words, designs without an unexposed reference and without baseline adjustment would substantially overstate how specifically BRS predicts recovery after adversity.

### Polygenic scores validation

To assess whether our PGS depended on the choice of LDpred2 model, we compared LDpred2-inf with LDpred2-auto in the three settings used in this study. LDpred2 is a Bayesian method that re-estimates variant effect sizes from GWAS summary statistics, accounting for linkage disequilibrium (LD) between variants using an LD reference panel. Both models assume a point-normal prior on variant effects, but they differ in how the genetic architecture is specified. LDpred2-inf treats all variants as causal (*p* = 1) and requires the SNP heritability as a fixed input, which yields a closed-form solution, whereas LDpred2-auto estimates both the proportion of causal variants (*p*) and the SNP heritability (h²) directly from the data by Gibbs sampling, without a separate validation sample for tuning. We derived and validated three sets of PGS: the first two were GS PGS derived from the GWAS discovery cohort (N = 159,575) and from the GWAS-for-PGS cohort (N = 130,726). The third was UKB PGS derived from the GWAS-for-PGS cohort (Fig. S3).

LDpred2-auto converged well in all three settings. All 30 chains passed quality control in each setting, and the chains agreed closely on the SNP heritability (Fig. S3a). The median per-chain h² was 6.96%, 7.80% and 7.92% in the three settings. The between-chain standard deviation was small: 0.12, 0.18 and 0.35 percentage points, or 1.7–4.4% of the estimate.

LDpred2-inf and LDpred2-auto produced highly similar PGS. Their scores were strongly correlated within each target cohort (r = 0.91, 0.90 and 0.97; Fig. S3b), and both showed a similar monotonic gradient of BRS across PGS percentile groups (Fig. S3c). Their effect sizes on BRS were also comparable (Fig. S3d):

- GS, PGS from the GWAS discovery cohort: standardized β = 0.080 (95% CI 0.059–0.101) for LDpred2-inf and 0.080 (0.059–0.101) for LDpred2-auto.
- GS, PGS from the GWAS-for-PGS cohort: 0.077 (0.056–0.098) and 0.076 (0.055–0.097).
- UKB, PGS from the GWAS-for-PGS cohort: 0.067 (0.055–0.078) and 0.062 (0.050–0.073).

Paired bootstrap comparisons (5,000 resamples) showed no difference between the two models in GS (Δβ = −0.0001, *P* = 0.97; Δβ = −0.001, *P* = 0.78). In UKB, LDpred2-auto was slightly lower (Δβ = −0.005, 95% CI −0.008 to −0.002, *P* = 0.0004). Thus, LDpred2-auto did not improve prediction in any setting.

Because the two models performed similarly, we used LDpred2-inf for all main analyses. Given the SNP heritability, LDpred2-inf has a closed-form solution, so it gives identical scores each time it is run. LDpred2-auto, by contrast, relies on MCMC sampling from random starting values and on criteria for which chains to keep, so its scores can vary slightly between runs. LDpred2-inf therefore provides the more reproducible basis for the downstream imaging-genetics analyses.

**Fig. S3.**
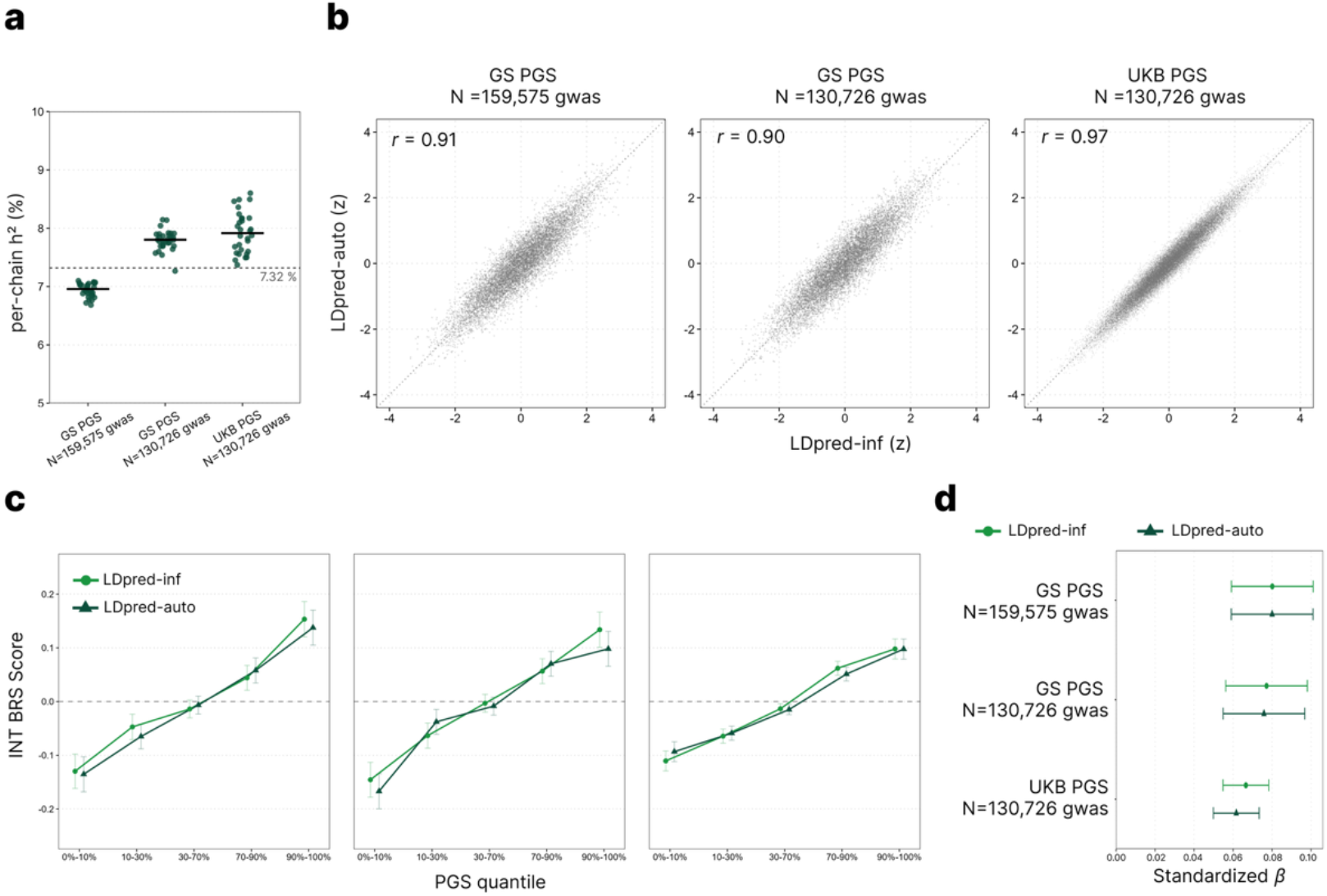
Comparison of LDpred2-inf and LDpred2-auto polygenic scores for BRS. PGS were derived with LDpred2 under the infinitesimal model (LDpred2-inf) and the automatic model (LDpred2-auto). They were calculated from GWAS of the GWAS discovery cohort (N = 159,575) or the GWAS-for-PGS cohort (N = 130,726; Fig. S1b) and applied to the GS PGS replication cohort (N = 8,702) and the UKB imaging-genetics discovery cohort (N = 27,729). The UKB imaging-genetics discovery cohort was not part of the 130,726-participant GWAS. **a,** SNP heritability (h²) estimated by LDpred2-auto for each GWAS–target combination. Each point is the estimate from one of 30 chains, each started from a different initial value of the causal-variant proportion *p*. Black bars mark the median across chains, and the dashed line marks the LD score regression (LDSC). All 30 chains passed quality control in each setting (see Methods). **b,** Agreement between LDpred2-inf and LDpred2-auto PGS (both standardized) within each target cohort. The dotted line mark’s identity, and r is the Pearson correlation coefficient. **c,** Mean INT BRS score by PGS percentile group (0–10%, 10–30%, 30–70%, 70–90% and 90–100%) for LDpred2-inf and LDpred2-auto, with panels in the same order as **b**. **d,** Standardized β of the PGS from linear regression of INT BRS on PGS, with both variables standardized and no covariates. Error bars represent ±1 standard error in **c** and 95% confidence intervals in **d**.

### Genetic enrichment analysis on human brain

The Linnarsson atlas used in Fig. 2e is derived from mouse. To examine whether the cell-type pattern extends to human brain tissue, we repeated the FUMA cell type analysis with the Allen human brain atlas. The two atlases differ in an important respect. The Linnarsson atlas spans the whole mouse nervous system, so the specificity of a cell type reflects both its class and its region; telencephalic glutamatergic neurons are contrasted with, among others, hindbrain, spinal cord and peripheral cell types. The human analysis also included fewer cell types and was based on nuclear rather than whole-cell transcriptomes. We therefore expected weaker signals and treated this analysis as a test of whether the relative ordering of cell classes was preserved, not as an independent replication.

No cell type survived FDR correction in either region. Three cell types reached nominal significance, all of them MTG glutamatergic neurons: Exc L6 FEZF2 SCUBE1 (P = 0.0052), Exc L5–6 SLC17A7 IL15 (P = 0.020) and Exc L3–5 RORB FILIP1L (P = 0.035) (Fig. S4). Beyond these three, the ordering of cell classes in the MTG resembled that in the mouse telencephalon: nine of the ten top-ranked MTG cell types were glutamatergic; the exception was oligodendrocyte precursor cells (P = 0.080). In the LGN, no cell type reached nominal significance (smallest P = 0.077, oligodendrocyte precursor cells), and the three thalamic relay neuron types showed no enrichment (all P ≥ 0.35). These results are consistent with, but weaker than, the mouse findings and should be regarded as supporting rather than confirmatory evidence.

**Supplementary Fig. 4.**
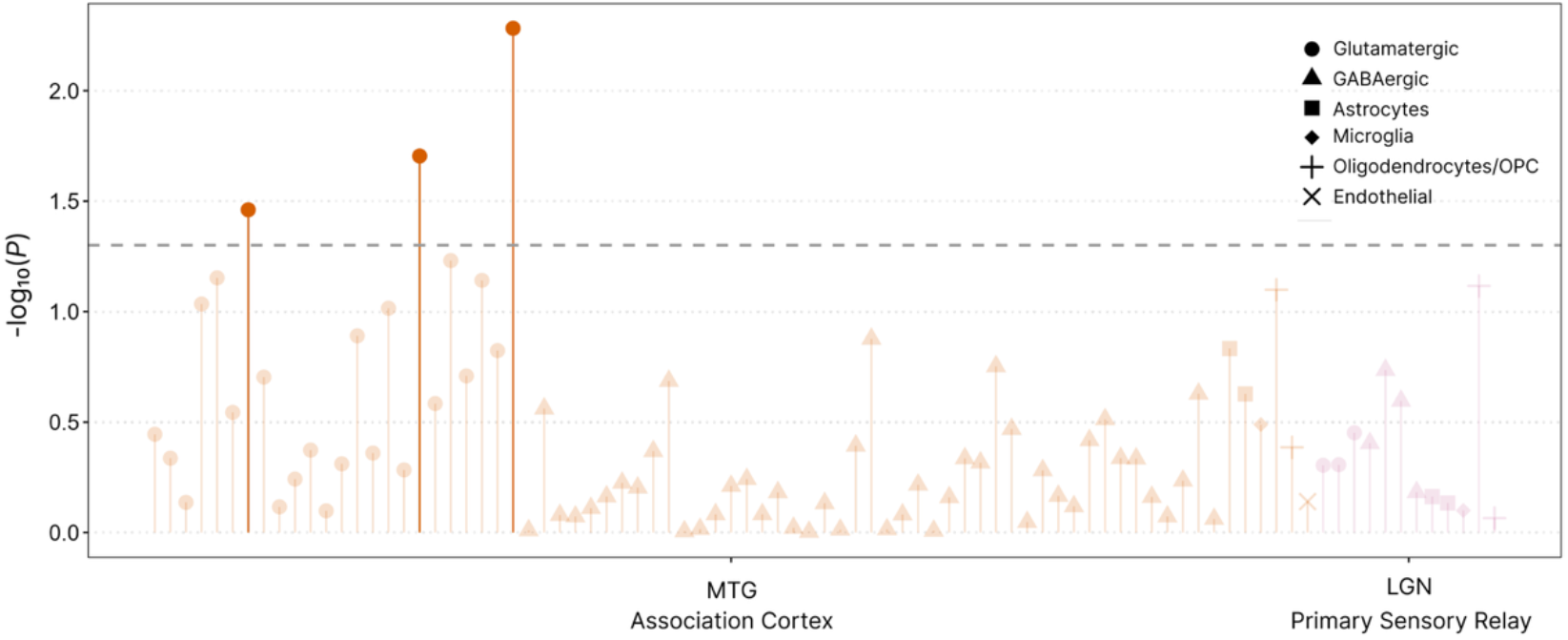
Cell-type-specific enrichment of BRS in the Allen human brain atlas. FUMA cell type analysis, of the BRS GWAS from the UK Biobank discovery cohort (N = 159,575) across 75 cell types of the middle temporal gyrus (MTG; association cortex) and 12 cell types of the lateral geniculate nucleus (LGN; thalamic primary sensory relay). Colors indicate brain regions (orange, MTG; pink, LGN), and shapes indicate major cell classes. Highlighted symbols denote three MTG glutamatergic cell types that reached nominal significance, none of which survived FDR correction within each dataset (all P_FDR ≥ 0.39). Dashed line, nominal P = 0.05

### Multivariate rsfMRI predictive model

To examine whether out-of-sample prediction depended on the modeling approach, we compared the three network-specific models trained in UKB (Fig. S5). All three models are linear combinations of standardized FCs but differ in how the weights are derived. With one component, the PLS weight of each FC is proportional to its edge-wise correlation with residualized INT BRS. CPM uses the same edge-wise correlations but coarsens them: FCs nominally associated with INT BRS (*P* < 0.01) receive a weight of +1 or −1 according to the sign of their association, and all other FCs receive a weight of zero. Ridge regression instead estimates all weights jointly, so the weight of an FC reflects its association conditional on the other, highly correlated FCs.

Both CPM and PLS derive their edge weights from the same quantity, the correlation between each FC and INT BRS: with standardized data, the one-component PLS weight of each FC is proportional to this correlation, whereas CPM retains only FCs whose correlation reaches nominal significance (*P* < 0.01) and weights them equally. CPM therefore selected the FCs with the largest absolute PLS weights, and the two models differed only in how the selected FCs were weighted. All selected FCs were positively associated with INT BRS (Fig. S5b). Ridge weights were moderately correlated with PLS weights (*r* = 0.75–0.87), but ridge assigned the opposite sign to 23 of 45, 5 of 17 and 30 of 57 FCs, including FCs that were individually positively associated with INT BRS (Fig. S5b). Because neighboring FCs are strongly correlated, these opposite-signed weights likely reflect contrasts between FCs rather than their shared signal, and ridge scores in GS diverged from PLS scores (*r* = 0.52–0.77; Fig. S5a). Together, these results indicate that the UKB association is carried by many weak, same-signed FC effects, which CPM and PLS aggregate in similar ways. Ridge, by contrast, gave opposite-signed weights to FCs that were highly correlated with each other. Its scores depended on small differences between similar FCs rather than on their shared signal. We suggest such differences are more likely to reflect features specific to the UKB sample, which may explain why ridge generalized less well to GS.

**Fig. S5.**
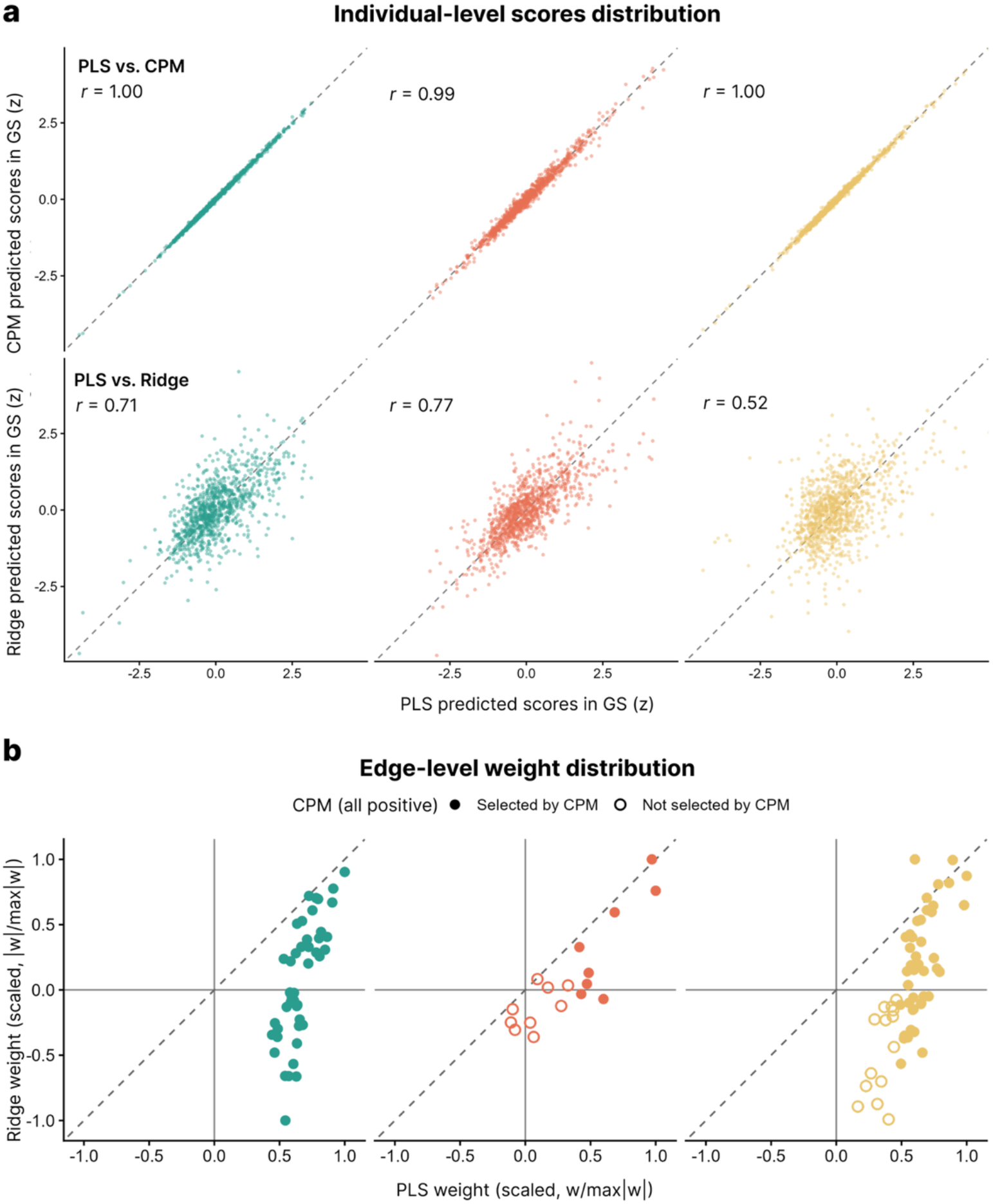
Comparison of the three UKB-trained predictive models. Network-specific models (CPM, PLS with one component and ridge regression) were trained in the UKB imaging-genetics discovery cohort (N = 27,729) after residualizing INT BRS and each FC on age, mean-centered age squared, sex, log head motion and log tSNR, and were applied without refitting to the GS imaging replication cohort (N = 1,060). Columns correspond to H1– H3 (colors as in Fig. 3). **a,** Individual-level predicted scores in GS. Each point represents one participant; scores are standardized within each model. Top row, PLS versus CPM; bottom row, PLS versus ridge. The dashed line marks identity, and r is the Pearson correlation coefficient. **b,** Edge-level model weights. Each point represents one FC. Weights are scaled by the maximum absolute weight of each model within each network (w/max|w|). Filled circles mark FCs selected by CPM (P < 0.01 for the association with residualized INT BRS; all selected FCs were positively associated), and open circles mark FCs not selected. The dashed line marks identity.

### Replication of edge-wise association

As a complement to the network-level prediction analysis (Fig. 3d), we compared edge-wise associations between cohorts. Because single-edge effects were small (UKB partial r = 0.012–0.052) and the GS sample was modest (N = 1,060), the GS replication cohort had limited power to detect individual edges. Accordingly, no GS edge survived FDR correction (13 of 45 H1 edges reached nominal P < 0.05), and we evaluated directional consistency rather than the significance of individual edges.

GS estimates matched the UKB direction for 44 of 45 H1 edges (the exception, vmPFC–vmPFC, had r = −0.001 in GS), 13 of 17 H2 edges and 54 of 57 H3 edges. A binomial test treats each edge as an independent trial that matches the UKB direction with a probability of 0.5, and would therefore suggest overwhelming significance (e.g., H1, P = 1.3 × 10⁻¹²). However, edges within a network are correlated, so under the null they tend to shift in the same direction together; this widens the null distribution of sign concordance and makes high concordance far more likely by chance than the binomial test assumes. We therefore generated null distributions by permuting BRS within each GS imaging site, which preserves the correlations among edges. The resulting null is wide and flat when most edges share the same UKB direction (H1 and H3), because correlated shifts then move all edges toward or away from concordance together, and approaches the binomial shape when UKB directions are mixed (H2), because such shifts then partly cancel out (Fig. S6a). Under these nulls, sign concordance was significant for H1 (P = 0.042) and H3 (P = 0.029), as was the sign-aligned mean partial correlation (H1, mean r = 0.048, P = 0.011; H3, mean r = 0.044, P = 0.009; Fig. S6b), whereas neither measure was significant for H2 (P ≥ 0.11). Only the permutation results are reported in the main text, because the binomial null ignores the dependence among edges and therefore overstates significance.

Because H1 showed the most robust replication, we further examined its edges by subnetwork and compared their directions with those most frequently reported in prior literature (Fig. S6c). In both cohorts, edges within and between the CEN, SN and DMN were uniformly positively associated with BRS. This agreed with the positive associations commonly reported within the CEN and between the CEN and DMN, extended to connections for which prior findings are mixed (within the DMN and between the SN and the CEN or DMN), and contrasted with the negative association commonly reported within the SN (see Discussion).

**Fig. S6.**
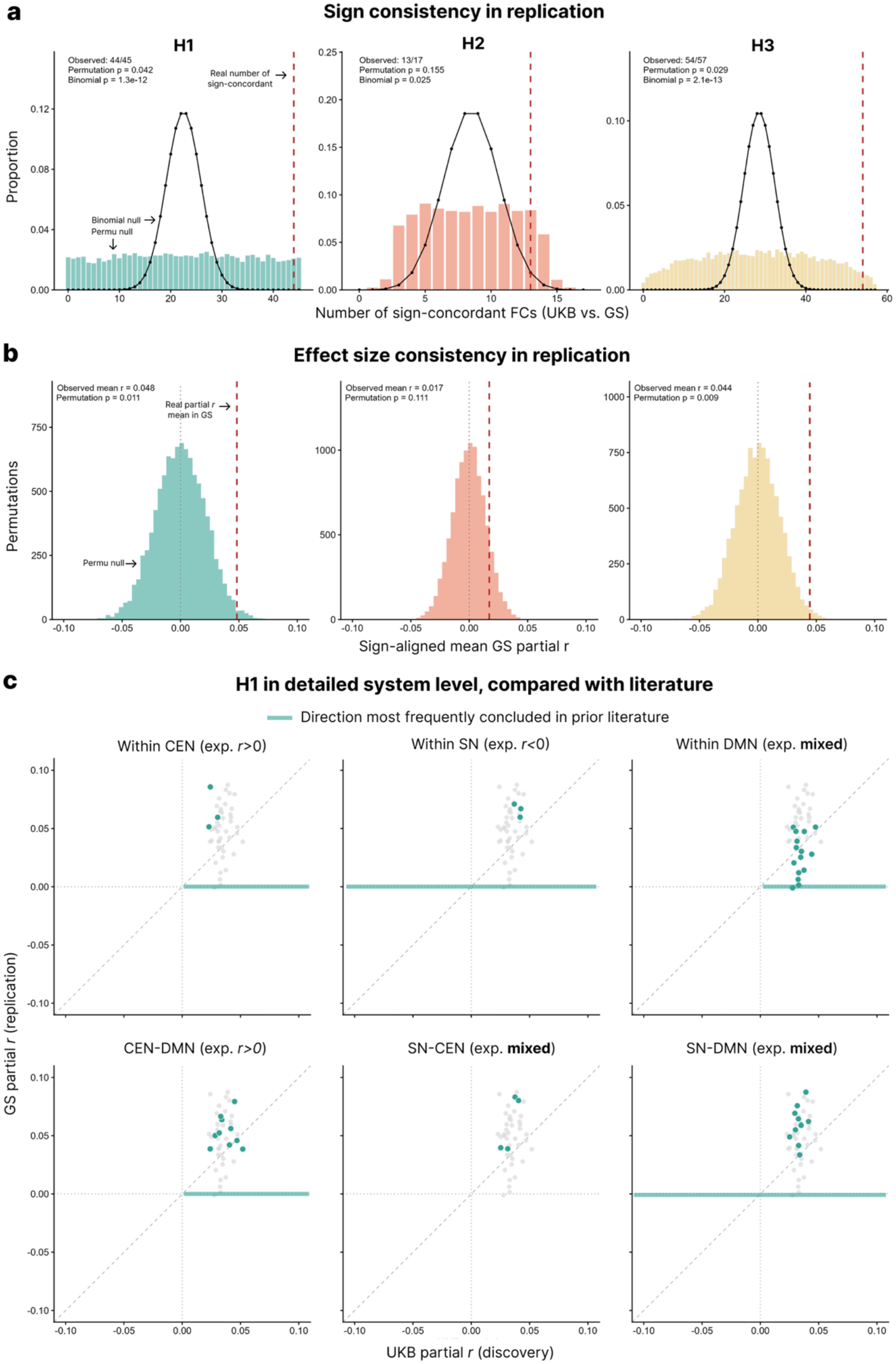
Consistency of edge-wise associations between the UKB discovery and GS replication cohorts. Edge-wise partial correlations were estimated in UKB (N = 27,729) and GS (N = 1,060) as in Fig. 3b, e. Null distributions were generated by permuting INT BRS scores within each GS imaging site 10,000 times and re-estimating all edge-wise partial correlations with the same covariates, which preserves the correlation structure among FCs. **a,** Sign consistency. Number of FCs whose partial correlations had the same sign in UKB and GS under the permutation null (bars), compared with the Binomial (n, 0.5) distribution that assumes independent FCs (line). The red dashed line marks the observed number of sign-concordant FCs. **b,** Effect-size consistency. Permutation null distribution of the sign-aligned mean GS partial correlation (the mean of sign[r_UKB] × r_GS across FCs). The red dashed line marks the observed value, and the dotted line marks zero. **c,** H1 FCs by subnetwork. UKB versus GS edge-wise partial correlations for FCs within (CEN, SN, DMN) and between (CEN–DMN, SN–CEN, SN–DMN) the three subnetworks (colored points), shown against all H1 FCs (grey points). Panel titles indicate the direction of association most frequently reported in prior literature (horizontal bars). The dashed line marks identity.

### Replication of edge-wise mediation

Single-edge indirect effects in UKB were small (standardized a × b = 3.2–8.2 × 10⁻⁴), and the GS imaging-genetics replication cohort (N = 997) had limited power to detect them. Accordingly, no GS FC reached FDR significance. To quantify the expected precision at this sample size, we drew 5,000 random UKB subsamples of 997 participants without replacement and repeated the edge-wise mediation analysis in each subsample. In these subsamples, the power to detect the indirect effects of the 13 FCs significant in UKB was below 1%, and no subsample yielded any FDR-significant FC. We therefore evaluated whether the indirect effects replicated in direction at the network level rather than for individual edges.

GS indirect effects matched the UKB direction for 42 of 45 FCs in H1, 12 of 17 in H2 and 27 of 57 in H3. As for the edge-wise associations (Fig. S6), a binomial test that treats FCs as independent overstates significance (e.g., H1, P = 4.3 × 10⁻¹⁰). We therefore generated null distributions by permuting the PGS within each GS imaging site, which disrupts the PGS–FC (a) path while preserving the correlations among FCs and their associations with BRS. Under this null, sign concordance (P = 0.047) and the sign-aligned mean standardized indirect effect (2.1 × 10⁻³, P = 0.022) were significant for H1 (Fig. S7a, b), whereas neither measure was significant for H2 (P = 0.072 and P = 0.070) or H3 (P = 0.55 and P = 0.57).

The GS estimates were on average 4.2-fold larger than those in UKB (range 0.2–8.0 across the 13 FCs significant in UKB). Two observations indicate that this difference reflects the limited precision of the smaller cohort rather than a failure to replicate. First, the UKB estimate fell within the GS 95% confidence interval for all 45 H1 FCs (Fig. S7c, left). Second, the GS estimates fell within the distributions obtained from the UKB subsamples (44th–99th percentile; median, 89th), albeit toward their upper end (Fig. S7c, right). Thus, the mediation of polygenic resilience through H1 connectivity was consistent across cohorts in direction, whereas estimating the indirect effects of individual connections requires samples of UKB scale.

**Fig. S7.**
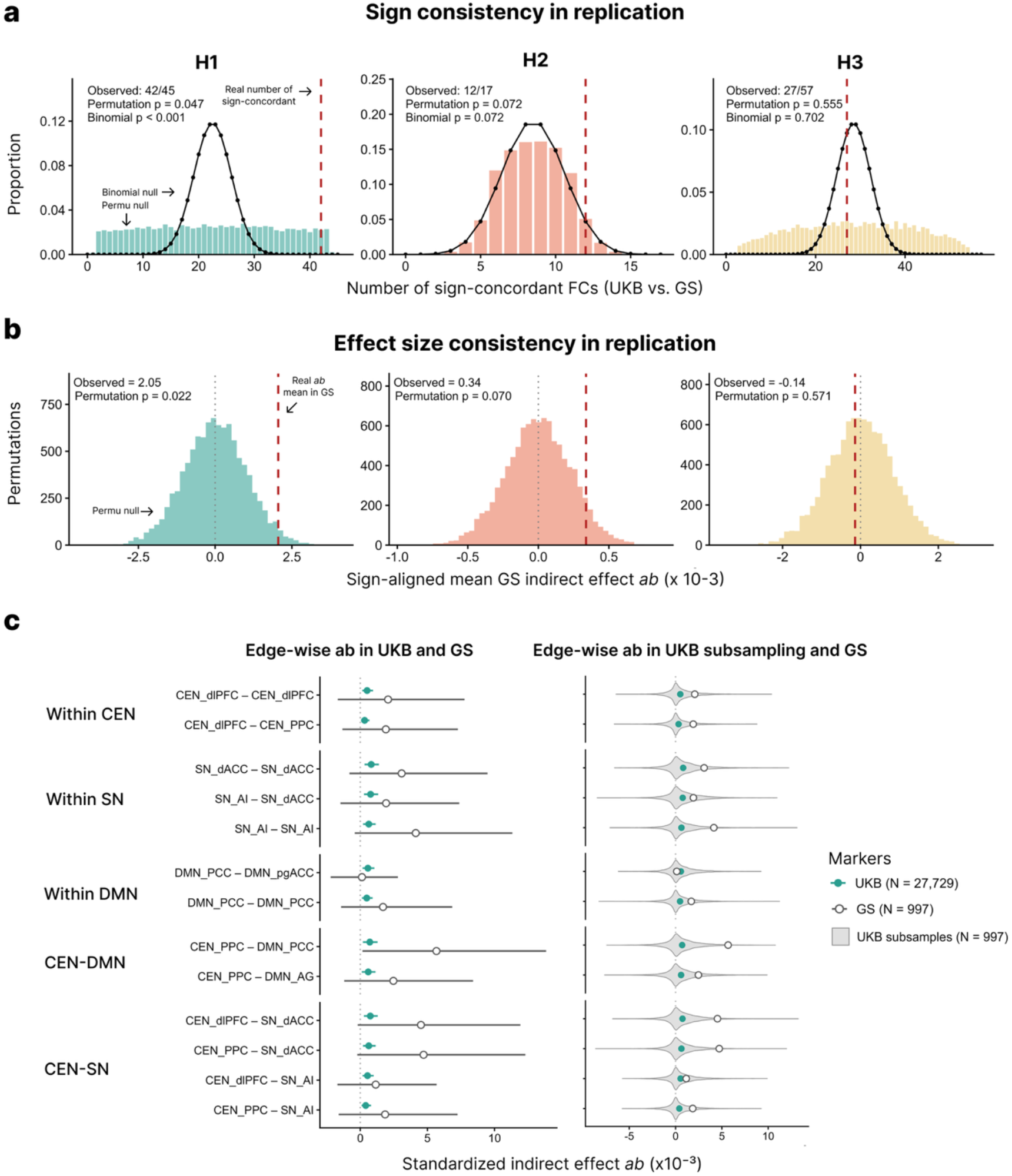
Replication of edge-wise mediation in the GS imaging-genetics replication cohort. **a,** Sign consistency. For each network, the number of FCs whose indirect effect in GS (N = 997) had the same sign as in UKB (dashed red line). Bars show the null distribution from 10,000 permutations of the PGS within imaging site, which preserves the correlations among FCs; the black line shows the binomial distribution (n, 0.5), which assumes independent FCs. **b,** Effect size consistency. Mean standardized indirect effect in GS after aligning each FC to the sign of its UKB estimate (dashed red line), compared with the null distribution from the same permutations. **c,** Standardized indirect effects of the 13 H1 FCs with FDR-significant indirect effects in UKB, grouped by subnetwork. Left, estimates in UKB (filled circles; N = 27,729) and GS (open circles; N = 997) with 95% Monte Carlo confidence intervals. Right, distributions of the estimates across 5,000 random UKB subsamples of 997 participants (violins), with the full-sample UKB (filled) and GS (open) estimates overlaid. All models were adjusted for age, age squared, sex, log head motion and log tSNR, with imaging site additionally included in GS.

### Replication of system integration

To assess whether the SSI findings generalized beyond UKB, we focused on the seven SSIs significantly associated with BRS in UKB (Fig. 4e, f) and tested them in GS using the same SSI definitions and models, with scanning site as an additional covariate. Unlike the edge-wise analyses, these SSIs were too few and too strongly interdependent for network-level concordance tests: they are derived from the same within- and between-network FC blocks, and each one-to-triple SSI equals the mean of the two corresponding pairwise SSIs. We therefore evaluated each SSI individually, testing associations in the direction observed in UKB (one-sided) and comparing the magnitudes of the indirect effects between cohorts.

All seven SSI–BRS associations had the same direction in GS as in UKB (N = 1,060; Fig. S8, left). Integration of the DMN replicated most clearly: DMN integration into the triple network (r = −0.056, one-sided P = 0.034) and with the SN (r = −0.062, one-sided P = 0.023) were significant, with GS estimates comparable to or larger than those in UKB (r = −0.033 and −0.029). Associations involving CEN integration were weaker in GS (e.g., CEN into the triple network, r = −0.019, one-sided P = 0.27, vs r = −0.047 in UKB). The power of the GS cohort to detect associations of the UKB magnitudes was 9–33%.

For mediation (N = 997; Fig. S8, right), the standardized indirect effects through CEN integration into the triple network (0.52 × 10⁻³) and with the SN (0.61 × 10⁻³) closely matched those in UKB (0.63 × 10⁻³ and 0.57 × 10⁻³), but their confidence intervals were wide and included zero (P ≥ 0.71). The pattern across SSIs also resembled that in UKB: although all seven SSIs were associated with BRS, the indirect effects in UKB were confined to CEN integration into the triple network and with the SN (shaded rows), whereas CEN integration with the DMN, which showed the strongest pairwise association with BRS, did not mediate the PGS effect. Thus, the SSI associations were directionally consistent across cohorts, and the indirect effects through CEN integration were similar in magnitude, although the replication cohort lacked the power to detect effects of this size.

**Fig. S8.**
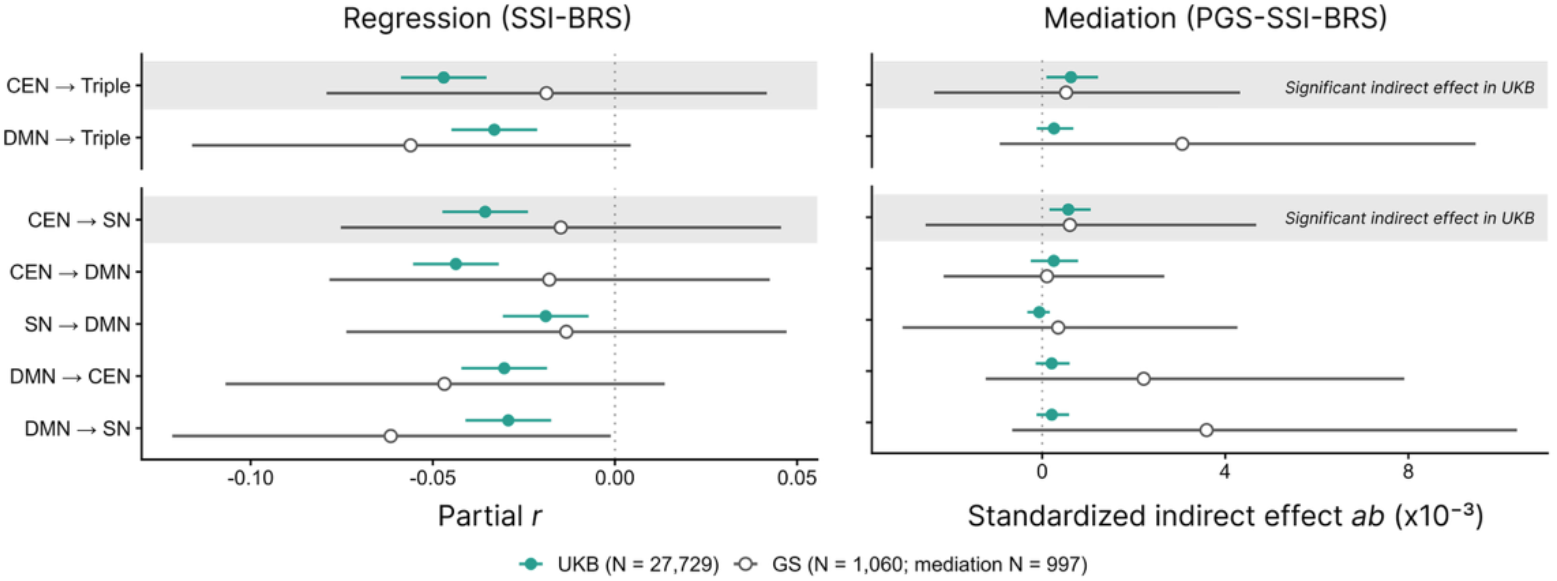
System-level SSI associations and mediation in UKB and GS. The seven SSIs significantly associated with INT BRS in UKB, shown for one-to-triple (top) and pairwise (bottom) SSIs; X → Y denotes the segregation of network X from Y (or from the triple network). Left, partial correlations between SSI and INT BRS; right, completely standardized indirect effects (a × b) of the PGS on INT BRS through SSI. Error bars show 95% confidence intervals. Shaded rows indicate SSIs with significant indirect effects in UKB (P < 0.05). Models were adjusted for covariates.

### Replication of childhood environmental effect

We examined whether the CEQ findings generalized to the GS imaging-genetics replication cohort. CEQ in GS was derived from the 25 items of the CTQ-SF; as in UKB, higher scores reflect a more supportive, non-maltreating early-life environment. Of the 997 GS participants, 971 had complete CTQ-SF data. All models were identical to those in UKB (N = 20,603), with imaging site as an additional covariate. Because the CTQ-SF (25 items) measures childhood experiences in more detail than the five-item Childhood Trauma Screener used in UKB, effect sizes involving CEQ are not directly comparable between cohorts, and we focused on whether the direction and significance of the associations replicated.

At the behavioral level, the GS results reproduced the additive pattern observed in UKB (Fig. S9a). The PGS was not associated with CEQ (UKB, β = 0.013, P = 0.055; GS, β = 0.047, P = 0.15), and CEQ and the PGS were each associated with BRS when modeled jointly (CEQ: UKB, β = 0.138, P = 4.4 × 10⁻⁹⁰; GS, β = 0.307, P = 6.5 × 10⁻²³; PGS: UKB, β = 0.066, P = 6.9 × 10⁻²²; GS, β = 0.086, P = 0.005), without a significant interaction (UKB, β = 0.003, P = 0.65; GS, β = 0.034, P = 0.29).

At the network level, CEQ was associated with six SSIs in UKB (shaded rows in Fig. S9b): CEN and DMN integration into the triple network (r = −0.026 and −0.024) and both directions of the CEN–DMN and SN–DMN pairs (r = −0.029 to −0.016). All six associations replicated in GS in the same direction (r = −0.059 to −0.115, one-sided P ≤ 0.033), with larger effect sizes, consistent with the more detailed CEQ measure. In addition, CEQ in GS was associated with CEN integration with the SN (r = −0.100, P = 0.002), an association that did not reach significance in UKB (r = −0.013, P = 0.056). Thus, the link between CEQ and the DMN-centered integration features was robust across cohorts, whereas the separation between the genetically associated CEN–SN axis and the environmentally associated features was less distinct in GS. Associations with SN integration into the triple network and SN integration with the CEN were not significant in either cohort.

**Fig. S9.**
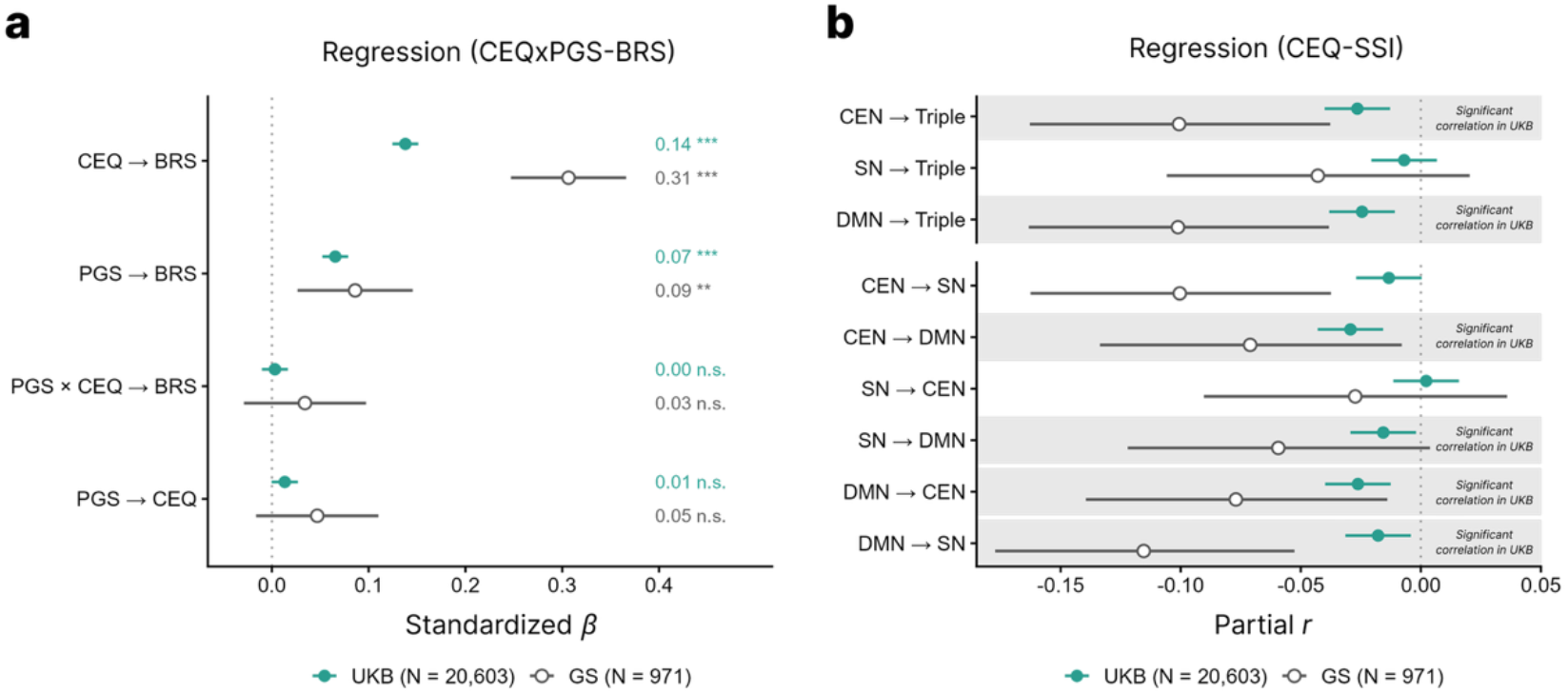
Replication of CEQ associations in the GS imaging-genetics replication cohort. Analyses included GS imaging-genetics replication cohort participants with complete CTQ-SF (N = 971), and UKB participants as in Fig. 5 (N = 20,603). Filled circles, UKB; open circles, GS; error bars, 95% confidence intervals. a, Standardized regression coefficients (β) of the PGS, CEQ and their interaction on BRS, and of the PGS on CEQ, estimated with the same models as in Fig. 5c, with imaging site additionally included in GS. Estimates and significance are shown on the right (**P < 0.01, ***P < 0.001; n.s., not significant). b, Partial correlations between CEQ and the nine SSIs, shown for one-to-triple (top) and pairwise (bottom) SSIs; X → Y denotes the segregation of network X from Y (or from the triple network). Shaded rows indicate SSIs significantly associated with CEQ in UKB (P < 0.05). Models were adjusted for covariates.

### MRI data processing in GS

The pipeline for deriving these FC matrices is designed to approximate the UKB approach as closely as possible. In GS, we process 3D T1-weighted, defaced data and resting-state BOLD data to obtain functional connectivity (FC) matrices based on the Glasser and Tian Scale II atlases.

The GS dataset comprises two MRI scanning protocols, referred to as the Aberdeen and Dundee protocols^5^: for T1-weighted imaging, the Dundee protocol closely matches the UKB protocol, using 1.0 mm isotropic voxels and a 256 × 256 matrix with 208 slices, whereas Aberdeen has the same voxel size but a relatively smaller field of view, with a 240 × 240 matrix and 160 slices. T1 acquisition parameters were otherwise broadly comparable across sites: Repetition Time (TR) was 2000 ms for UKB, 1968 ms for Aberdeen, and 1740 ms for Dundee, while Inversion Time (TI) was 880 ms for UKB, 1031 ms for Aberdeen, and 900 ms for Dundee. Greater differences were observed for resting-state fMRI (rsfMRI). Both Aberdeen and Dundee acquired data at a spatial resolution of 3.4 × 3.4 × 4.5 mm³ using a 64×64 matrix and 32 slices), whereas UKB employed a higher-resolution protocol with 2.4 mm isotropic voxels, an 88 × 88 matrix and 64 slices). Differences extended to temporal acquisition parameters. Aberdeen and Dundee utilized a TR of 1560 ms, acquiring 195 timepoints over 5 minutes and 4 seconds. In contrast, UKB employed highly accelerated scanning techniques with a TR of 735 ms, enabling the collection of 490 timepoints in 6 minutes and 10 seconds. Further differences were evident in image contrast parameters: Aberdeen and Dundee used a shorter Echo Time (TE=26ms and 22 ms, respectively) and a 70° flip angle, while UKB used a longer TE (39 ms) and a smaller flip angle (52°) to accommodate its shorter TR.

### 1.1 Preprocessing of T1 and BOLD data with FSL

In UKB, rsfMRI BOLD time series are processed with the Melodic pipeline in FSL (FMRIB Software Library)^6^, using individual preprocessed T1 images, brain-extracted T1 images, linear and non-linear warp of T1 to MNI152 space, and binary mask of white matter as structural references^7^. T1 structural images are first preprocessed by gradient distortion correction (GDC) and FOV reduction of non-brain regions. Linear and non-linear registration to MNI152 space is utilized to the preprocessed T1 images to extract the warps of T1 to MNI152 space. Using the inverse of this MNI152 alignment warp, a standard-space brain mask is transformed into native T1 space and applied to the T1 image to generate a skull-stripped, brain-extracted T1^7^. The brain-extracted T1 is further used to segment tissue types with the FAST tool, yielding a binary mask of white matter. To process the T1 structural images in the GS dataset, we implemented the open-source UK Biobank pipeline (https://git.fmrib.ox.ac.uk/falmagro/UK_biobank_pipeline_v_1). Notably, the gradient distortion correction (GDC) step was bypassed, as this correction was already applied on-scanner during the GS data acquisition.

In the processing of rsfMRI BOLD data, UKB resamples the data by simultaneously applying GDC transformation, head motion correction and EPI unwarping, where the unwarping process relies on Boundary-Based Registration (BBR) to align the functional data to the subject’s high-resolution T1 structural image^7^. Subsequent steps include standardization of overall 4D time series intensity via a global multiplicative constant, followed by a high-pass temporal filtering, which is implemented through a Gaussian-weighted least-squares straight line fitting (sigma = 50.0s) ^7^. Artifacts are than removed by using ICA+FIX^7^. For the rsfMRI BOLD data in the GS dataset, we adapted the same open-source UKB functional pipeline described above with a few necessary modifications. First, given the relatively longer TR in the GS cohort (1560 ms compared to 735 ms in UKB), we incorporated a slice-timing correction step to account for temporal offsets during volume acquisition. Consistent with the structural processing, GDC was omitted. Furthermore, because neither AP/PA reverse phase-encoded scans nor B0 field maps were acquired, susceptibility distortion correction (EPI unwarping) was not performed. Instead, the alignment of functional data to the T1 structural images relied exclusively on BBR. For data denoising, the pre-trained ICA+FIX classifier utilized in the standard UKB pipeline was incompatible with our dataset due to discrepancies in rsfMRI acquisition parameters. Consequently, we implemented ICA-AROMA as an alternative approach for robust artifact removal^8^.

### 1.2 Surface segmentation of T1 with FastSurfer

While FSL provides robust volumetric tissue segmentation, mapping the Glasser cortical atlas requires precise surface-based representations (as detailed in the following section)^9^. To achieve this, T1-weighted images typically require processing via FreeSurfer. However, given the large computational cost of running the standard FreeSurfer pipeline on thousands of subjects, we implemented FastSurfer, a deep-learning alternative, to efficiently generate the necessary surface reconstructions^10^.

### 1.3 Calculation of parcellated BOLD time series

After processing and quality control of T1 and BOLD data, BOLD time series for each subject is then parcellated to Glasser cortical atlas and Tian scale II atlas according to the pipeline used in UKB^9^. For the cortex, Glasser atlas, which is originally built on fsLR space template, was resampled to FreeSurfer’s fsaverage template, and then aligned to subjects’ native space (fsnative)^9^. These surface labels were mapped to the volumetric cortical ribbon by assigning each voxel the label of its nearest surface vertex^9^. For subcortical regions, the inverse of the pre-computed non-linear warp (which registers the subject’s T1 to MNI152 space) is applied to transform the Tian atlas from MNI152 space into the subject’s native T1 space^9^. After mapping the atlas to the subject’s native T1 space, the volumetric parcellation is resampled to the preprocessed fMRI voxel grid^9^. For each parcel, BOLD time series is extracted by averaging the signal across all voxels within that region^9^. We follow the same procedures of parcellation mentioned above in GS. Parcel-level FC was computed as the Pearson correlation between the time series. ROI-level FC was then defined as the mean Fisher z value across all parcel pairs connecting two anatomical ROIs; for connections within a single ROI, only the unique pairs of distinct parcels were averaged. Missing values were imputed separately within each imaging site (Aberdeen and Dundee) using k-nearest-neighbour imputation (k = 20) in which each missing value was replaced by the median of the 20 participants with the most similar ROI-level FC profiles. Two rsfMRI data-quality measures, mean absolute head motion and temporal signal-to-noise ratio (tSNR), were included as covariates in all analyses. Both measures were derived to approximate the UKB IDP definitions. Mean absolute head motion was computed from the MCFLIRT rigid-body transformation matrices as the root-mean-square displacement within an 80-mm-radius sphere of each volume relative to the first volume, averaged across volumes, with the temporal-mean image as reference. tSNR was computed from the ICA-AROMA-denoised parcel time series as the median, across parcels, of the temporal mean divided by the temporal standard deviation.

**Fig. S10.**
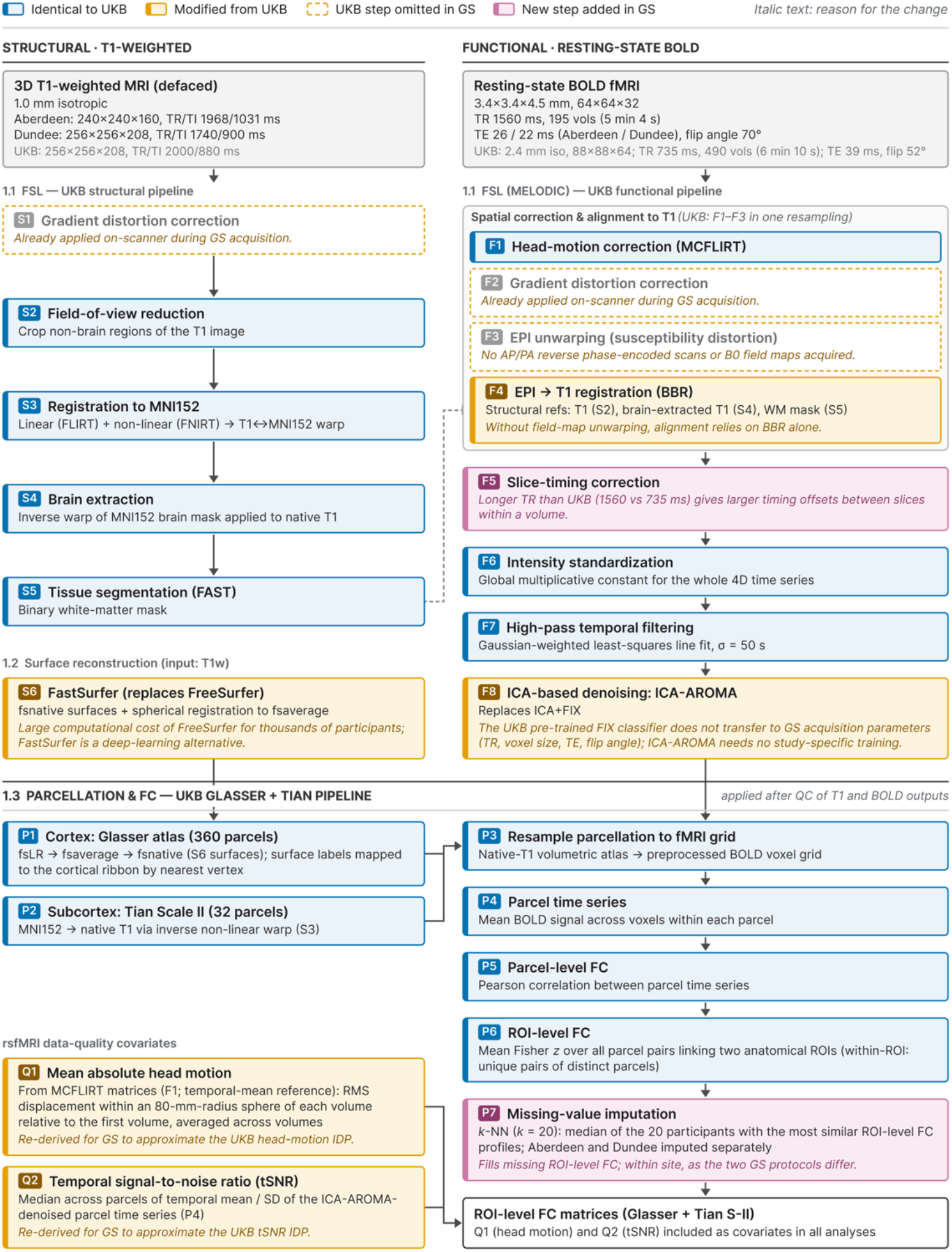
MRI processing in GS relative to the UKB pipeline. Structural (left) and resting-state functional (right) data from both GS protocols (Aberdeen and Dundee) were processed with the open-source UKB pipeline, followed by Glasser and Tian Scale II parcellation and ROI-level FC estimation; numbered headings refer to the corresponding sections of the Supplementary Methods. Colors indicate correspondence with UKB: identical (blue), modified (orange, solid outline), UKB steps omitted in GS (orange, dashed outline) and steps added in GS (purple); italic text gives the reason for each deviation. Grey boxes summarize acquisition parameters, with UKB values in grey text. Step labels: S, structural processing; F, functional processing; P, parcellation and FC estimation; Q, rsfMRI data-quality covariates. Solid arrows indicate data flow; the dashed arrow indicates structural references used for BBR. GDC, gradient distortion correction; BBR, boundary-based registration; IDP, imaging-derived phenotype; WM, white matter.

### Brain hypothesis

The following paragraphs introduce our criteria for mapping Glasser and Tian S2 parcels onto hypothesis-driven ROIs and systems.

**Fig. S11.**
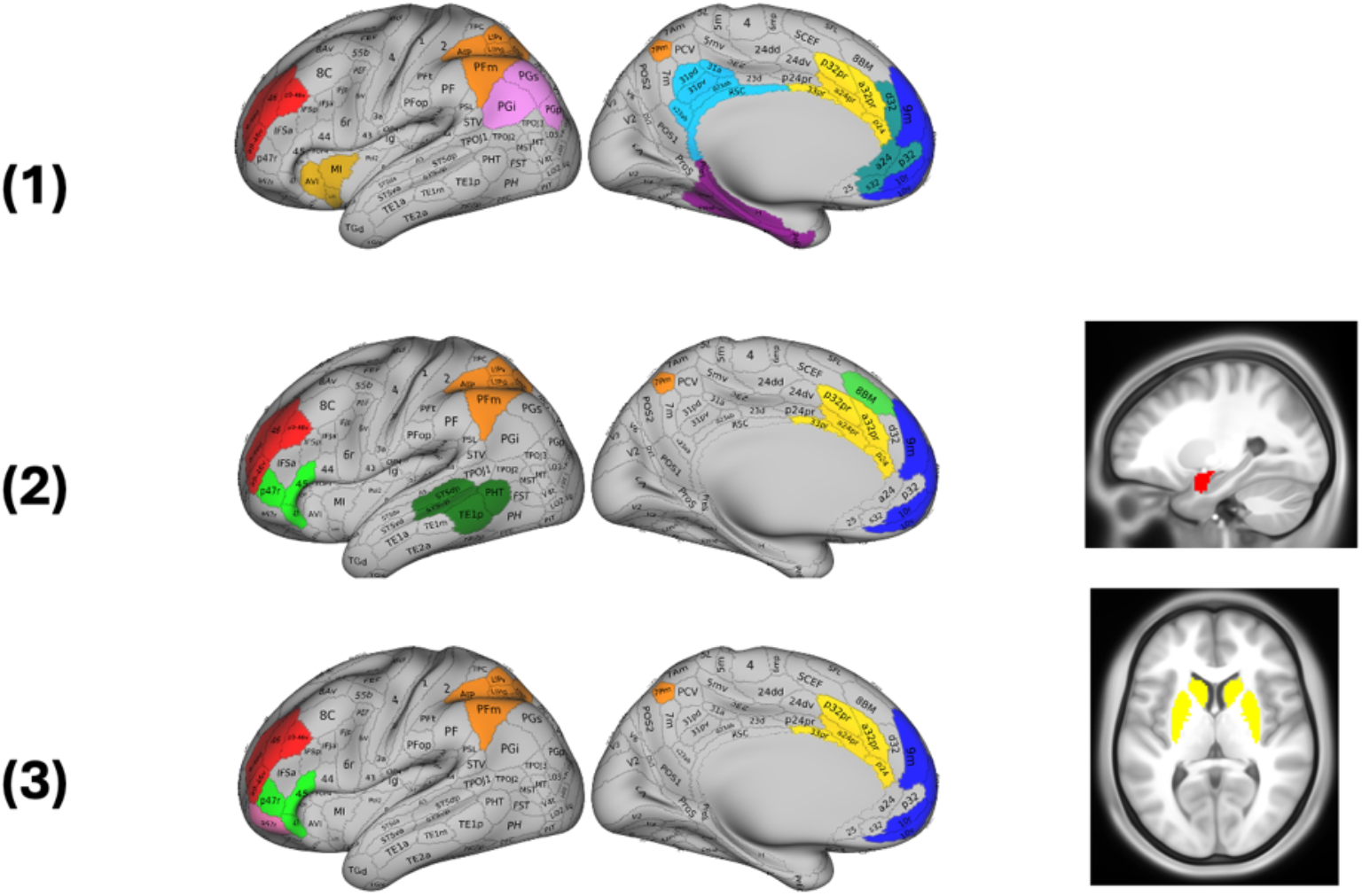
Anatomical definitions of the ROIs in the three hypothesized brain systems. Cortical ROIs were defined from the Glasser (HCP-MMP1.0) parcellation and subcortical ROIs from the Tian Scale II atlas; ROIs were defined bilaterally and are shown on the lateral and medial surfaces of the left hemisphere. **(1)** Large-scale flexibility network (H1): the CEN (dlPFC: 46, 9-46d, a9-46v, p9-46v; PPC: LIPv, LIPd, AIP, MIP, 7Pm, PFm, IP1, IP2), SN (dACC: a32pr, p32pr, a24pr, p24, 33pr; AI: AVI, AAIC, MI) and DMN (vmPFC: 10r, 10v, 10d, 9m; pgACC: s32, a24, p32, d32; PCC: RSC, v23ab, d23ab, 31pv, 31pd, 31a; MTL: H, PreS, EC, PeEc, PHA1–3; AG: PGi, PGs, PGp). **(2)** Cortico-amygdala regulatory network (H2): the dlPFC, vlPFC (45, 47l, p47r), dmPFC (8BM), PPC, dACC, vmPFC and lateral temporal cortex (LTC: STSdp, STSvp, TE1p, PHT), together with the lateral and medial amygdala (sagittal slice). **(3)** Cortico-striatal valuation network (H3): the dlPFC, vlPFC, OFC (a47r, p47r), PPC, dACC and vmPFC, together with the striatum (core and shell of the nucleus accumbens and anterior and posterior caudate and putamen; axial slice). Colors denote ROIs as in Fig. 3a. The rationale for each ROI is described in the text.

**(1) Attention and cognitive control system (triple network model)**

i. According to Menon 2011, representative regions of CEN include the dorsolateral prefrontal cortex (dlPFC) and the posterior parietal cortex (PPC)^11^. The dlPFC is subdivided into 13 cortical areas: 8C, 8Av, i6-8, s6-8, SFL, 8BL, 9p, 9a, 8Ad, p9-46v, a9-46v, 46, and 9-46d according to the HCP-MMP1.0 parcellation proposed by Glasser et al. 2016 (Supplementary Neuroanatomical Results, p.76)^12^. However, this represents a relatively broad anatomical delineation of the dlPFC. In the present study, we selected four parcels—9-46d, 46, a9-46v, and p9-46v—as representative dlPFC regions for the CEN, as these parcels correspond to the central portion of the dlPFC in the Glasser parcellation (Supplementary Neuroanatomical Results, p.79). The latter one, PPC, is anatomically defined as the parietal association cortex encompassing the superior parietal cortex/lobule (SPL), inferior parietal cortex/lobule (IPL), and intraparietal sulcus (IPS), following by the classical framework of Mesulam 1998^13^. Within the HCP-MMP1.0 atlas, the superior parietal cortex together with the medial bank of the intraparietal sulcus (IPS) is subdivided into ten parcels: LIPv, LIPd, VIP, AIP, MIP, 7PC, 7AL, 7Am, 7PL, and 7Pm (Supplementary Neuroanatomical Results, p.52). Inspection of the working-memory contrast (2BK–0BK) provided in the Glasser supplementary material indicates that only a subset of these regions shows robust task-positive responses. Specifically, VIP, 7PC, 7AL, 7Am, and 7PL exhibit relatively weak activation in the 2BK–0BK contrast (Supplementary Neuroanatomical Results, p.52; Fig. 19) and were therefore excluded from the PPC definition of the CEN. Similarly, the inferior parietal cortex together with the lateral bank of the IPS is subdivided into ten parcels: PGp, PGs, PGi, PFm, PF, PFt, PFop, IP0, IP1, and IP2 (Supplementary Neuroanatomical Results, p.56). Among these, PGp, PGi, and PGs exhibit task-negative responses rather than task-positive activation, displaying a default-mode-like functional profile (Supplementary Neuroanatomical Results, p.56 & p.58). In addition, their anatomical locations are adjacent to the angular gyrus (AG; see Caspers et al., 2006)^14^, further supporting their classification as components of the default mode network rather than the CEN. We therefore excluded PGp, PGi, and PGs from the PPC definition. Furthermore, PF, PFt, PFop, and IP0 show relatively weak activation in the 2BK–0BK working-memory contrast (Supplementary Neuroanatomical Results, p.57; Fig. 20) and were also excluded. After applying these criteria, the remaining parcels PFm, IP1, and IP2 from the IPL/lateral IPS group, together with LIPv, LIPd, AIP, MIP, and 7Pm from the SPL/medial IPS group, were retained as the principal PPC nodes of the CEN. Together, both their functional response pattern and anatomical situation support classifying these regions as components of the DMN rather than the CEN. In summary, a total of 12 of 180 cortical parcels in the HCP-MMP1.0 atlas were classified as belonging to CEN.
ii. According to Menon 2011, DMN includes posterior cingulate cortex (PCC), medial prefrontal cortex (mPFC), medial temporal lobe (MTL) and angular gyrus (AG). For PCC, the anchor node of DMN, Glasser et al. 2016 subdivided 14 areas: DVT, ProS, POS1, POS2, RSC, v23ab, d23ab, 31pv, 31pd, 31a, 23d, 23c, PCV, and 7m (Supplementary Neuroanatomical Results, p.58)^12^. DVT and ProS were excluded because Glasser et al. describe them as transitional visual areas, with ProS showing strong functional connectivity with early visual cortex and DVT with higher dorsal-stream visual cortex (Supplementary Neuroanatomical Results, p. 59-60). POS1 and POS2 were not treated as core PCC nodes because the parietal-occipital sulcus region is characterized as a transitional belt between early visual cortex and posterior cingulate association cortex (Supplementary Neuroanatomical Results, p.59). Area 7m was also excluded from the core PCC definition because, although it shows connectivity with task-negative parietal regions, its task-activation profile is not consistent with a stable DMN core (Supplementary Neuroanatomical Results, p.60). As a result, regions of RSC, v23ab, d23ab, 31pv, 31pd and 31a are defined as our PCC nodes for DMN. In functional network parcellations, the mPFC node of the default mode network (DMN) is typically centered on ventromedial prefrontal cortex (vmPFC), whereas dorsomedial prefrontal cortex (dmPFC) is often treated as functionally distinct (Andrews-Hanna et al., 2010^15^; Margulies et al., 2016^16^). By the work of Rolls et al. 2022, within the HCP-MMP1.0 atlas, vmPFC regions consist of 10r, 10v, 10d and 9m, while pregenual anterior cingulate cortex (pgACC) regions consist of s32, a24, p24, p32, d32, as these regions show strong functional and effective connectivity to PCC^17^. However, we did not include p24 in the pgACC set because, from an anatomical perspective, p24 is located closer to the mid-cingulate / dorsal ACC belt within the area 24 complex. Instead, p24 was included as part of the dACC region in the subsequent section. As a result, we defined the remaining regions as our DMN-anchored mPFC ROI. For MTL, we use HCP-MMP1.0 atlas defined 7 areas in medial temporal cortex: H, PreS, Ec, PeEc, PHA1, PHA2, and PHA3 (Supplementary Neuroanatomical Results, p.44) ^12^. For AG, we have claimed PGp, PGi, and PGs in the HCP-MMP1.0 atlas should be classified into AG as discussed in the former part. According to Caspers, 2006 ^14^, AG is discerned into an anterior portion (PGa) and posterior portion (PGp). When mapping this scheme onto the HCP-MMP1.0 atlas, PGa corresponds primarily to PGi together with PGs, whereas PGp corresponds directly to PGp (Supplementary Neuroanatomical Results, p.56 & p.58) ^12^. Accordingly, we conclude that PGp, PGi, and PGs collectively provide an appropriate representation of the angular gyrus for DMN labeling in the HCP-MMP1.0 atlas.
iii. According to Menon 2011, SN-anchored regions are dorsal anterior cingulate cortex (dACC) and anterior insula (AI). dACC anatomically corresponds to midcingulate cortex (MCC) ^18^, and MCC is parcellated into 24dd and 24dv in HCP-MMP1.0 atlas. However, these two regions are more motor-related output regions (Supplementary Neuroanatomical Results, p.26-p.28), which does not align with the concepts of dACC in SN. According to intracranial stimulation on cingulate cortex reported by Oane et al. 2020, regions a32pr, p32pr, a24pr, p24, and 33pr labeled as anterior cingulate in HCP-MMP1.0 atlas are anatomically located on anterior MCC (aMCC), and they proposed the aMCC as a region that couples affective signals with motor systems involved in goal-directed actions by emotional processing of negative stimuli (fear, anger, panic)^19^. As a result, we define 5 areas of aMCC a32pr, p32pr, a24pr, p24, and 33pr as our dACC for SN. AI is clearly delineated in HCP-MMP1.0 atlas by 3 areas: AVI, AAIC, and MI (Supplementary Neuroanatomical Results, p.42-p.43)^12^.

**(2) Emotion regulation system (cognitive reappraisal)**

The amygdala is the most commonly modulated region in studies of reappraisal. According to Ochsner, Silvers, and Buhle (2012), reappraisal consistently recruits domain-general cognitive control regions, including the dlPFC, vlPFC, and dmPFC, together with the PPC, which supports attention and working memory processes. The dACC may monitor whether the ongoing emotional response is changing in the intended direction during reappraisal. In addition, the vmPFC is thought to integrate affective valuation signals and may function as a mediating node linking prefrontal control regions with the amygdala^20^.

Based on this framework, we selected FC sets that reflect the hypothesized interactions between cognitive control systems and affect systems during reappraisal. Specifically, we examined connectivity between domain-general cognitive control regions (dlPFC, vlPFC, dmPFC, PPC and dACC) and the valuation-related vmPFC with the amygdala, as well as intra-amygdala connectivity to capture potential changes in affective processing within the amygdala.

The dlPFC and PPC were previously defined as components of the CEN and we therefore adopted their Glasser atlas ROIs from the CEN definition. The vmPFC was defined in the DMN section and included Glasser parcels 10r, 10v, 10d, 9m. The dACC was previously defined as part of the SN and was represented by the Glasser parcels a32pr, p32pr, a24pr, p24, 33pr. The amygdala was defined using the Tian atlas, which separates the amygdala into medial and lateral subdivisions; both subdivisions were included in the analyses. The vlPFC is chosen as 45, 47I, p47r. These regions show activation in the relational-match and story contrasts in the HCP task maps and are anatomically located within the inferior frontal cortex (Supplementary Neuroanatomical Results, p.73 Fig. 24). As for dmPFC, according to Assem et al. (2020), area 8BM shows a multiple-demand activation pattern and lies within the dorsomedial prefrontal cortex. Therefore, we used 8BM as a proxy for the dmPFC in the reappraisal network^21^.

Another line of research suggests that activation in the vmPFC is not consistently observed across empirical studies^22^. Instead, activation in the lateral temporal cortex—associated with sensory semantic processing—appears to be a more consistent pattern. These findings suggest that reappraisal may also operate through modulation of semantic representations^22^.

Prior work has also identified several lateral temporal cortex regions as key nodes of the semantic network. Milton et al. (2021) demonstrated that multiple lateral temporal parcels in the HCP-MMP atlas—including STSdp, STSvp, TE1p, and PHT—show consistent activation across semantic categorization tasks. These regions correspond to the posterior superior temporal sulcus and middle temporal cortex, which are widely implicated in semantic representation and conceptual processing. Therefore, we included these parcels (STSdp, STSvp, TE1p, and PHT) as regions of interest representing the lateral temporal semantic cortex. As a result, we examined their functional connectivity with the amygdala to assess whether semantic representations in lateral temporal cortex may influence affective responses during reappraisal.

**(3) Reward and decision system**

The review of Wolfram Schultz^23^ has suggested that reward and decision signals are encoded across a distributed cortico–striatal system. In addition to dopaminergic neurons, value-related signals have been observed in multiple cortical regions, including the OFC, vmPFC, vlPFC, dlPFC, anterior ACC, and PPC, as well as in the striatum and amygdala. These regions form key components of the reward valuation system through cortico–striatal and striato–amygdala interactions. However, the dopaminergic nuclei that generate reward prediction error signals (e.g., the substantia nigra pars compacta and ventral tegmental area) are relatively small midbrain structures and are not included in the Tian atlas used in the present study. Therefore, interactions involving these dopaminergic nuclei with the striatum and cortical regions were not examined in the current analysis^23^.

Accordingly, we operationalized the reward valuation system using functional connectivity (FC) sets reflecting interactions among these regions. Specifically, we examined FC between cortical valuation regions (OFC, vmPFC, vlPFC, dlPFC, anterior ACC, and PPC) and the striatum, FC between the striatum and the amygdala, and intra-striatal connectivity. These connectivity sets were used to represent the cortico–striatal and striato–amygdala components of the reward valuation system.

We retained the definitions of vmPFC, vlPFC, dlPFC, and PPC parcellations in the HCP-MMP atlas as described in the previous two sections. The striatum was defined using the Tian S2 atlas and included shell NAc, core NAc, anterior caudate, posterior caudate, anterior putamen, and posterior putamen.

For the OFC, the naming system of parcellation follows the scheme proposed by Öngür et al. (2003)^24^. In that framework, areas 47/12r, 47/12l, 47/12m, and 47/12s correspond to the lateral orbital cortex. Therefore, we focused on the corresponding parcels in the HCP-MMP atlas, namely a47r, 47m, 47s, and p47r. Furthermore, because only a47r and p47r show visible activation in the relational-match and gambling tasks (Supplementary Neuroanatomical Results, p.69, Fig. 23), we selected these three parcels as proxies for reward- and decision-related OFC.

Regarding the anterior ACC mentioned by Wolfram Schultz^23^, this term refers to an anatomically broad region. After reviewing the literature cited by Schultz, we found that these studies primarily focused on the dorsal ACC (dACC)^25^, which is predefined in our salience network (SN). Therefore, we used the parcels a32pr, p32pr, a24pr, p24, and 33pr as proxies for dACC.

## References

1. Kalisch, R. et al. The resilience framework as a strategy to combat stress-related disorders. Nat. Hum. Behav. 1, 784–790 (2017).

2. Kalisch, R., Russo, S. J. & Müller, M. B. Neurobiology and systems biology of stress resilience. Physiol. Rev. 104, 1205–1263 (2024).

3. Ungar, M. & Theron, L. Resilience and mental health: how multisystemic processes contribute to positive outcomes. Lancet Psychiatry 7, 441–448 (2020).

4. Thakkar, K. N. et al. Moving from risk to resilience in psychosis research. Nat. Rev. Psychol. 2, 537–555 (2023).

5. Choi, K. W., Stein, M. B., Dunn, E. C., Koenen, K. C. & Smoller, J. W. Genomics and psychological resilience: a research agenda. Mol. Psychiatry 24, 1770–1778 (2019).

6. Nestler, E. J. & Russo, S. J. Neurobiological basis of stress resilience. Neuron 112, 1911–1929 (2024).

7. Russo, S. J., Murrough, J. W., Han, M.-H., Charney, D. S. & Nestler, E. J. Neurobiology of resilience. Nat. Neurosci. 15, 1475–1484 (2012).

8. Visscher, P. M. et al. 10 Years of GWAS Discovery: Biology, Function, and Translation. Am. J. Hum. Genet. 101, 5–22 (2017).

9. Marek, S. et al. Reproducible brain-wide association studies require thousands of individuals. Nature 603, 654–660 (2022).

10. Bonanno, G. A., Chen, S., Bagrodia, R. & Galatzer-Levy, I. R. Resilience and Disaster: Flexible Adaptation in the Face of Uncertain Threat. Annu. Rev. Psychol. 75, 573–599 (2024).

11. Wolke, D. et al. A systematic review of conceptualizations and statistical methods in longitudinal studies of resilience. Nat. Ment. Health 3, 1088–1099 (2025).

12. Oshio, A., Taku, K., Hirano, M. & Saeed, G. Resilience and Big Five personality traits: A meta-analysis. Personal. Individ. Differ. 127, 54–60 (2018).

13. Cornelis, M. C. et al. The brief resilience scale: a genome-wide association study in the UK Biobank. BMC Med. 23, 520 (2025).

14. Norbury, A., Seeley, S. H., Perez-Rodriguez, M. M. & Feder, A. Functional neuroimaging of resilience to trauma: convergent evidence and challenges for future research. Psychol. Med. 53, 3293–3305 (2023).

15. Tabibnia, G. An affective neuroscience model of boosting resilience in adults. Neurosci. Biobehav. Rev. 115, 321–350 (2020).

16. Misaki, M. et al. Connectome-wide investigation of altered resting-state functional connectivity in war veterans with and without posttraumatic stress disorder. NeuroImage Clin. 17, 285–296 (2018).

17. Shi, L., Sun, J., Wei, D. & Qiu, J. Recover from the adversity: functional connectivity basis of psychological resilience. Neuropsychologia 122, 20–27 (2019).

18. Sripada, R. K. et al. Neural Dysregulation in Posttraumatic Stress Disorder: Evidence for Disrupted Equilibrium Between Salience and Default Mode Brain Networks. Psychosom. Med. 74, 904–911 (2012).

19. Engert, V., Blasberg, J. U., Köhne, S., Strauss, B. & Rosendahl, J. Resilience and personality as predictors of the biological stress load during the first wave of the Covid-19 pandemic in Germany. Transl. Psychiatry 11, 443 (2021).

20. Platania, S., Gruttadauria, S. V. & Morando, M. Dispositional Resilience as Mediator in Psychological Stress on Healthcare Workers: A Multi-Group Analysis of Frontline and Non-Frontline Workers. Eur. J. Investig. Health Psychol. Educ. 12, 1285–1299 (2022).

21. Semkovska, M., et al. Modelling the Relationship Between Resilience to Depression and Recent Stressful Life Events in University Students. Psychol. Rep. 127, 1561–1587 (2024).

22. Köhne, S., Engert, V. & Rosendahl, J. Stability of resilience in times of the COVID-19 pandemic. Personal. Ment. Health 17, 55–66 (2023).

23. Engert, V., Blasberg, J. U., Köhne, S., Strauss, B. & Rosendahl, J. Resilience and personality as predictors of the biological stress load during the first wave of the Covid-19 pandemic in Germany. Transl. Psychiatry 11, 443 (2021).

24. Troy, A. S. et al. Psychological Resilience: An Affect-Regulation Framework. Annu. Rev. Psychol. 74, 547–576 (2023).

25. Menon, V. Large-scale brain networks and psychopathology: a unifying triple network model. Trends Cogn. Sci. 15, 483–506 (2011).

26. Miller, G. E. et al. Functional connectivity in central executive network protects youth against cardiometabolic risks linked with neighborhood violence. Proc. Natl. Acad. Sci. 115, 12063–12068 (2018).

27. Chan, A. et al. Trauma and the default mode network: review and exploratory study. Front. Behav. Neurosci. 18, 1499408 (2024).

28. Rakesh, D., Allen, N. B. & Whittle, S. Longitudinal changes in within-salience network functional connectivity mediate the relationship between childhood abuse and neglect, and mental health during adolescence. Psychol. Med. 53, 1552–1564 (2023).

29. Zhang, L., Rakesh, D., Cropley, V. & Whittle, S. Neurobiological correlates of resilience during childhood and adolescence – A systematic review. Clin. Psychol. Rev. 105, 102333 (2023).

30. Kilpatrick, L. A. et al. Sex commonalities and differences in the relationship between resilient personality and the intrinsic connectivity of the salience and default mode networks. Biol. Psychol. 112, 107–115 (2015).

31. Shao, R., Lau, W. K. W., Leung, M.-K. & Lee, T. M. C. Subgenual anterior cingulate-insula resting-state connectivity as a neural correlate to trait and state stress resilience. Brain Cogn. 124, 73–81 (2018).

32. Tai, A. P. L., Leung, M.-K., Geng, X. & Lau, W. K. W. Conceptualizing psychological resilience through resting-state functional MRI in a mentally healthy population: a systematic review. Front. Behav. Neurosci. 17, 1175064 (2023).

33. Sripada, R. K. et al. Neural Dysregulation in Posttraumatic Stress Disorder: Evidence for Disrupted Equilibrium Between Salience and Default Mode Brain Networks. Psychosom. Med. 74, 904–911 (2012).

34. Bolsinger, J., Seifritz, E., Kleim, B. & Manoliu, A. Neuroimaging Correlates of Resilience to Traumatic Events—A Comprehensive Review. Front. Psychiatry 9, 693 (2018).

35. Minikel, E. V., Painter, J. L., Dong, C. C. & Nelson, M. R. Refining the impact of genetic evidence on clinical success. Nature 629, 624–629 (2024).

36. Wagner, A. Distributed robustness versus redundancy as causes of mutational robustness. BioEssays 27, 176–188 (2005).

37. Edelman, G. M. & Gally, J. A. Degeneracy and complexity in biological systems. Proc. Natl. Acad. Sci. 98, 13763–13768 (2001).

38. Smith, B. W. et al. The brief resilience scale: Assessing the ability to bounce back. Int. J. Behav. Med. 15, 194–200 (2008).

39. Davis, K. A. S. et al. The UK Biobank Mental Health Enhancement 2022: Methods and Results. Preprint at 10.1101/2024.11.21.24317700 (2024).

40. Habota, T. et al. Cohort profile for the STratifying Resilience and Depression Longitudinally (STRADL) study: A depression-focused investigation of Generation Scotland, using detailed clinical, cognitive, and neuroimaging assessments [version 2; peer review: 2 approved]. Wellcome Open Res. (2021).

41. Loh, P.-R., Kichaev, G., Gazal, S., Schoech, A. P. & Price, A. L. Mixed-model association for biobank-scale datasets. Nat. Genet. 50, 906–908 (2018).

42. Chang, C. C. et al. Second-generation PLINK: rising to the challenge of larger and richer datasets. Gigascience 4, s13742-015-0047–8 (2015).

43. Kerr, S. M. et al. Electronic health record and genome-wide genetic data in Generation Scotland participants. Wellcome Open Res. 2, 85 (2017).

44. Alfaro-Almagro, F. et al. Image processing and Quality Control for the first 10,000 brain imaging datasets from UK Biobank. NeuroImage 166, 400–424 (2018).

45. Mansour L., S., Di Biase, M. A., Smith, R. E., Zalesky, A. & Seguin, C. Connectomes for 40,000 UK Biobank participants: A multi-modal, multi-scale brain network resource. NeuroImage 283, 120407 (2023).

46. Glasser, M. F. et al. A multi-modal parcellation of human cerebral cortex. Nature 536, 171–178 (2016).

47. Tian, Y., Margulies, D. S., Breakspear, M. & Zalesky, A. Topographic organization of the human subcortex unveiled with functional connectivity gradients. Nat. Neurosci. 23, 1421–1432 (2020).

48. Hainmueller, J. Entropy Balancing for Causal Effects: A Multivariate Reweighting Method to Produce Balanced Samples in Observational Studies. Polit. Anal. 20, 25–46 (2012).

49. Loh, P.-R. et al. Efficient Bayesian mixed-model analysis increases association power in large cohorts. Nat. Genet. 47, 284–290 (2015).

50. Privé, F., Arbel, J. & Vilhjálmsson, B. J. LDpred2: better, faster, stronger. Bioinformatics 36, 5424–5431 (2021).

51. Privé, F., Albiñana, C., Arbel, J., Pasaniuc, B. & Vilhjálmsson, B. J. Inferring disease architecture and predictive ability with LDpred2-auto. Am. J. Hum. Genet. 110, 2042–2055 (2023).

52. Schizophrenia Working Group of the Psychiatric Genomics Consortium et al. LD Score regression distinguishes confounding from polygenicity in genome-wide association studies. Nat. Genet. 47, 291–295 (2015).

53. ReproGen Consortium et al. Partitioning heritability by functional annotation using genome-wide association summary statistics. Nat. Genet. 47, 1228–1235 (2015).

54. Weissbrod, O. et al. Functionally informed fine-mapping and polygenic localization of complex trait heritability. Nat. Genet. 52, 1355–1363 (2020).

55. De Leeuw, C. A., Mooij, J. M., Heskes, T. & Posthuma, D. MAGMA: Generalized Gene-Set Analysis of GWAS Data. PLOS Comput. Biol. 11, e1004219 (2015).

56. Wu, T. et al. clusterProfiler 4.0: A universal enrichment tool for interpreting omics data. The Innovation 2, 100141 (2021).

57. The Gene Ontology Consortium et al. The Gene Ontology knowledgebase in 2023. GENETICS 224, iyad031 (2023).

58. Watanabe, K., Umićević Mirkov, M., De Leeuw, C. A., Van Den Heuvel, M. P. & Posthuma, D. Genetic mapping of cell type specificity for complex traits. Nat. Commun. 10, 3222 (2019).

59. Zeisel, A. et al. Molecular Architecture of the Mouse Nervous System. Cell 174, 999–1014.e22 (2018).

60. Hodge, R. D. et al. Conserved cell types with divergent features in human versus mouse cortex. Nature 573, 61–68 (2019).

61. Preacher, K. J. & Selig, J. P. Advantages of Monte Carlo Confidence Intervals for Indirect Effects. Commun. Methods Meas. 6, 77–98 (2012).

## References

1. Hainmueller, J. Entropy Balancing for Causal Effects: A Multivariate Reweighting Method to Produce Balanced Samples in Observational Studies. Polit. Anal. 20, 25–46 (2012).

2. Kroenke, K., Spitzer, R. L., Williams, J. B. W. & Löwe, B. An Ultra-Brief Screening Scale for Anxiety and Depression: The PHQ–4. Psychosomatics 50, 613–621 (2009).

3. Kalisch, R. et al. The resilience framework as a strategy to combat stress-related disorders. Nat. Hum. Behav. 1, 784–790 (2017).

4. Bonanno, G. A., Chen, S., Bagrodia, R. & Galatzer-Levy, I. R. Resilience and Disaster: Flexible Adaptation in the Face of Uncertain Threat. Annu. Rev. Psychol. 75, 573–599 (2024).

5. Habota, T. et al. Cohort profile for the STratifying Resilience and Depression Longitudinally (STRADL) study: A depression-focused investigation of Generation Scotland, using detailed clinical, cognitive, and neuroimaging assessments [version 2; peer review: 2 approved]. Wellcome Open Res. (2021).

6. Jenkinson, M., Beckmann, C. F., Behrens, T. E. J., Woolrich, M. W. & Smith, S. M. FSL. NeuroImage 62, 782–790 (2012).

7. Alfaro-Almagro, F. et al. Image processing and Quality Control for the first 10,000 brain imaging datasets from UK Biobank. NeuroImage 166, 400–424 (2018).

8. Pruim, R. H. R. et al. ICA-AROMA: A robust ICA-based strategy for removing motion artifacts from fMRI data. NeuroImage 112, 267–277 (2015).

9. Mansour L., S., Di Biase, M. A., Smith, R. E., Zalesky, A. & Seguin, C. Connectomes for 40,000 UK Biobank participants: A multi-modal, multi-scale brain network resource. NeuroImage 283, 120407 (2023).

10. Henschel, L. et al. FastSurfer - A fast and accurate deep learning based neuroimaging pipeline. NeuroImage 219, 117012 (2020).

11. Menon, V. Large-scale brain networks and psychopathology: a unifying triple network model. Trends Cogn. Sci. 15, 483–506 (2011).

12. Glasser, M. F. et al. A multi-modal parcellation of human cerebral cortex. Nature 536, 171–178 (2016).

13. Mesulam, M. From sensation to cognition. Brain 121, 1013–1052 (1998).

14. Caspers, S. et al. The human inferior parietal cortex: Cytoarchitectonic parcellation and interindividual variability. NeuroImage 33, 430–448 (2006).

15. Andrews-Hanna, J. R., Reidler, J. S., Huang, C. & Buckner, R. L. Evidence for the Default Network’s Role in Spontaneous Cognition. J. Neurophysiol. 104, 322–335 (2010).

16. Margulies, D. S. et al. Situating the default-mode network along a principal gradient of macroscale cortical organization. Proc. Natl. Acad. Sci. 113, 12574–12579 (2016).

17. Rolls, E. T., Deco, G., Huang, C.-C. & Feng, J. The human orbitofrontal cortex, vmPFC, and anterior cingulate cortex effective connectome: emotion, memory, and action. Cereb. Cortex 33, 330–356 (2022).

18. Vogt, B. A., Nimchinsky, E. A., Vogt, L. J. & Hof, P. R. Human cingulate cortex: Surface features, flat maps, and cytoarchitecture. J. Comp. Neurol. 359, 490–506 (1995).

19. Oane, I. et al. Cingulate cortex function and multi-modal connectivity mapped using intracranial stimulation. NeuroImage 220, 117059 (2020).

20. Ochsner, K. N., Silvers, J. A. & Buhle, J. T. Functional imaging studies of emotion regulation: a synthetic review and evolving model of the cognitive control of emotion. Ann. N. Y. Acad. Sci. 1251, (2012).

21. Assem, M., Glasser, M. F., Van Essen, D. C. & Duncan, J. A Domain-General Cognitive Core Defined in Multimodally Parcellated Human Cortex. Cereb. Cortex 30, 4361–4380 (2020).

22. Buhle, J. T. et al. Cognitive Reappraisal of Emotion: A Meta-Analysis of Human Neuroimaging Studies. Cereb. Cortex 24, 2981–2990 (2014).

23. Schultz, W. Neuronal Reward and Decision Signals: From Theories to Data. Physiol. Rev. 95, 853–951 (2015).

24. Öngür, D., Ferry, A. T. & Price, J. L. Architectonic subdivision of the human orbital and medial prefrontal cortex. J. Comp. Neurol. 460, 425–449 (2003).

25. Seo, H. & Lee, D. Temporal Filtering of Reward Signals in the Dorsal Anterior Cingulate Cortex during a Mixed-Strategy Game. J. Neurosci. 27, 8366–8377 (2007).

